# DiffDose: Differentiable Programming for Personalized Dose-Regimen Optimal Control

**DOI:** 10.64898/2026.09.07.749974

**Authors:** Shayan Hajhashemi, Amin Emad, Morgan Craig

**Affiliations:** Department of Quantitative Life Sciences, McGill University, Montréal, QC, Canada; Mila, Quebec AI Institute, Montréal, QC, Canada; Sainte-Justine University Hospital Azrieli Research Centre, Montréal, QC, Canada; Department of Electrical and Computer Engineering, McGill University, Montréal, Canada; Rosalind and Morris Goodman Cancer Institute, Montréal, QC, Canada; Department of Mathematics and Statistics, Université de Montréal, Montréal, QC, Canada

**Keywords:** model-informed precision dosing, dosing regimen optimization, pharmacometrics, quantitative systems pharmacology, mechanistic PK/PD modeling, in silico clinical trials, virtual populations, differentiable programming, automatic differentiation, optimal control

## Abstract

Dose-regimen design requires choosing how much drug to give, when to give it, and how treatment should vary across patients. Mechanistic pharmacokinetic-pharmacodynamic (PK/PD) and quantitative systems pharmacology (QSP) models can predict treatment responses, but optimizing dosing inputs depends on model-specific sensitivity derivations or derivative-free search. Here, we introduce DiffDose, a differentiable programming framework for mechanistic open-loop dose-regimen optimization that uses automatic differentiation (AD) to handle clinically interpretable dose amounts and administration times as differentiable controls. We evaluate our method in three settings: fixed-schedule dose-amplitude optimization in OptiDose PK/PD benchmarks; dose-timing optimization in a chemotherapy-induced neutropenia model with state-dependent delay; and individualized mosunetuzumab dosing in a QSP virtual population. Across these examples, AD produced gradients consistent with references, reduced model-specific derivative work, and shortened benchmark time to solution. DiffDose thereby turns mechanistic PK/PD and QSP models from tools that evaluate prespecified regimens into gradient-based engines for individualized regimen design.

Code: https://github.com/Craig-Lab/DiffDose

## 1 Introduction

Drug therapy unfolds over time as a sequence of decisions shaped by disease progression. Dose-regimen design formalizes this problem: choosing how much drug to give^1,2^, when to give it^3,4^, and how those choices should vary across patients^5^. In practice, regimen selection is often empirical when drug modalities and disease indications allow for changes. Early-phase oncology uses dose-escalation rules to target acceptable toxicity rates^6^, while post-approval dosing may be refined through therapeutic drug monitoring or titration^7^. These approaches examine few schedules and offer limited mechanistic insight into changing disease and drug dynamics. These limitations are particularly important when response varies across patients or time, measured concentrations or biomarkers guide treatment, and exposure–response or pharmacokinetic/pharmacodynamic (PK/PD) relationships are non-flat during therapy; these scenarios may be caused by turnover delays, induction, target burden, and immunogenicity^8^.

Mathematical models, commonly formulated as systems of ordinary differential equations (ODEs), link administration events to evolving concentrations, effects, disease burden, biomarkers, and toxicity^9^. They range from PK/PD to mechanistic quantitative systems pharmacology (QSP) models. Once calibrated, they serve as in silico testbeds for candidate regimens and control-oriented decision support. Control theory frames this task as choosing a constrained input that minimizes an objective balancing efficacy, safety, adherence, and feasibility. Biomedical optimal control has a long history^10^, including early work on optimal drug regimens and saturation effects^11^ and applications in infectious disease and oncology^12–16^.

Classical optimal control often represents therapy as a continuous input, such as an infusion rate, and solves the resulting problem with Pontryagin or variational adjoint methods^17^. Many clinical regimens are discrete in time, magnitude, or both. Oral therapies, bolus injections, cycle-based dosing, dose holds, tablet strengths, and vial constraints are more naturally represented as finite events or jumps, motivating impulsive or hybrid ODEs^18–21^. Classical methods for optimal control of hybrid ODEs include direct multiple shooting and mixed-integer optimization^22,23^. Within pharmacometrics, Bachmann *et al*. recently introduced OptiDose, which to the best of our knowledge is the only published implementation that directly optimizes dose values for a fixed PK/PD schedule^24^. These workflows require model-specific considerations (e.g., sensitivity or adjoint derivations), increasing burden and reducing portability when the model, dosing event, objective, solver, or cohort changes^25–28^.

These constraints motivate a differentiable-programming approach to dose-regimen optimization. Hybrid jump conditions, saltation, and hidden discontinuities are treated in hybrid sensitivity theory^29–40^, more recently in Neural-ODEs^41–43^, and in optimal control^19,20,44,45^. Building on this literature, we introduce DiffDose, a differentiable programming framework that treats the mechanistic model, dosing events, numerical solver, quadrature, and clinical objective as a single differentiable computational graph. Automatic differentiation of this computational graph then propagates objective gradients to clinically interpretable dose amounts and administration times without a new hand-derived adjoint for each model^27,28^. We evaluate DiffDose across studies spanning fixed-time ODE dose-amplitude controls, dose-timing controls in a state-dependent delay differential equation (DDE)^46^, and patient-specific controls in a heterogeneous QSP virtual population (VPop)^47,48^. Across these settings, direct AD recovered hand-derived optima or finite-difference timing gradients and shortened time to solution in the benchmark comparisons. DiffDose therefore provides a reusable interface for turning mechanistic models from tools that compare prespecified regimens into tools that design patient-specific regimens, while retaining links among dosing decisions, model parameters, and outcomes.

## 2 Results

### 2.1 Differentiable hybrid control makes dose-regimen optimization modular

DiffDose uses automatic differentiation to separate dose-regimen design from model-specific derivative engineering. Each regimen is a finite set of controls connected to a mechanistic simulator (Figure 1). The same interface applies to PK/PD ODEs, DDEs, and QSP virtual populations, allowing the model or objective to change without rebuilding derivatives while keeping controls clinically readable.

**Figure 1.**
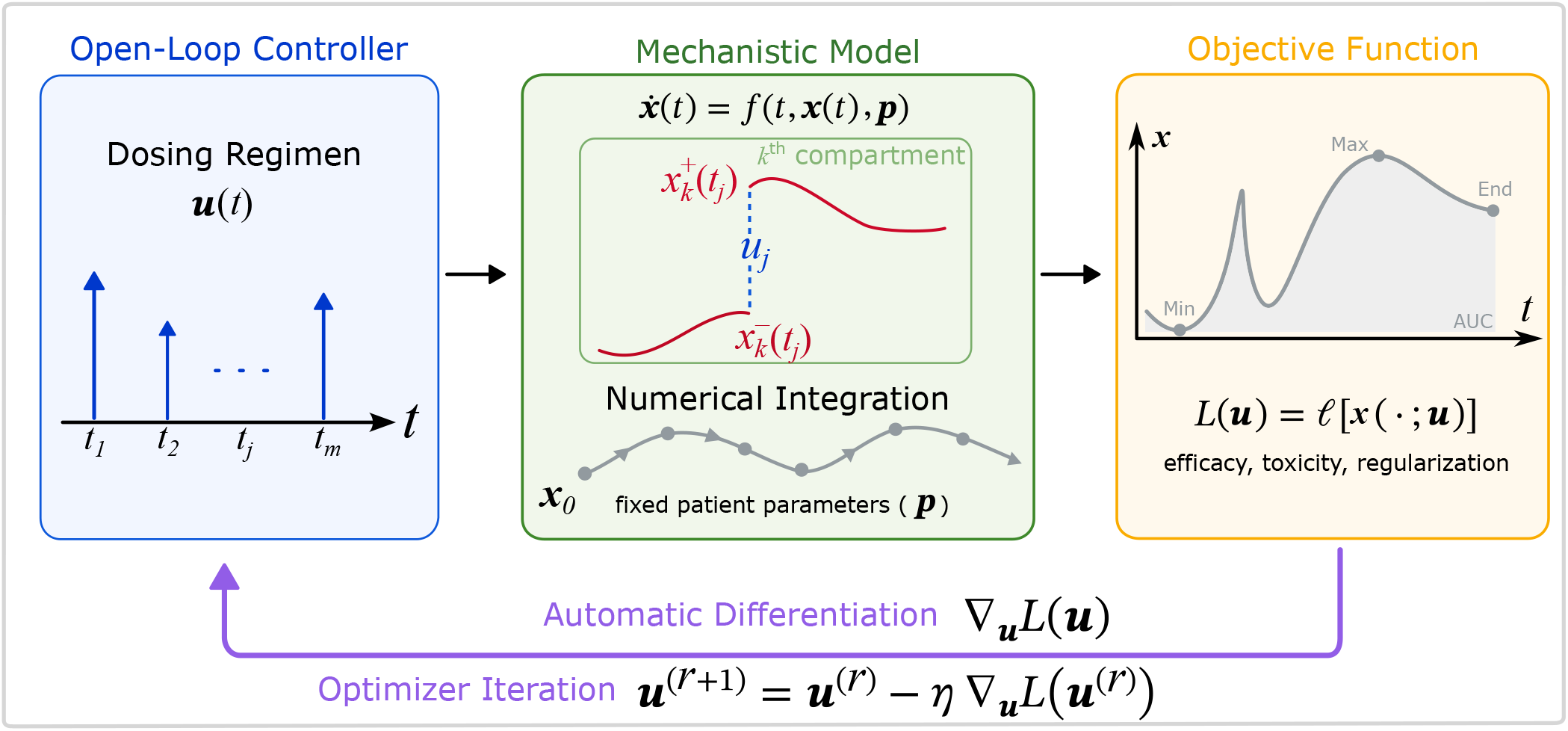
DiffDose uses differentiable programming to turn regimen design into an open-loop optimization problem. Schematic of input-output relationships of the differentiable computational graph. We start with a guess of the dosing regimen, **u**, and a given patient with parameters **p** and initial conditions **x**_0_ (blue box). We then numerically integrate the dynamics, f, until dose time t_j_ and update the dosing compartment with the corresponding dose amplitude, u_j_ (green box). We compute the loss functional, L, over the solution of the differential equation with the dosing events (orange box). Automatic differentiation formulas are used to obtain gradients that enable optimization of the open-loop dose regimen controls (purple arrow).

Consider a system of nonlinear ODEs, *f*, with states **x**(*t*) ∈ ℝ^*n*^ and parameters **p** ∈ ℝ^*q*^. For the derivation, assume one independent dose control *u*_*j*_ per fixed administration time *t*_*j*_, with **u** ∈ ℝ^*m*^. Each event is modeled by a reset map Φ_*j*_ : ℝ^*n*^ × ℝ → ℝ^*n*^, taking the pre-dose state and scalar dose to the post-dose state. Superscripts − and + denote left and right limits. Between dose administrations, the model evolves according to its usual dynamics. At administration times, the dose modifies the state through the reset map:

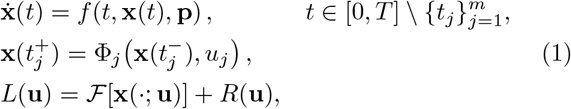

where ℱ is a scalar-valued trajectory functional encoding efficacy, safety, or exposure, and *R* is a direct dose penalty.

Instead of deriving model-specific sensitivities, DiffDose treats the simulator, event handling, quadrature, and loss as one differentiable program. For a perturbation **v** of the dose vector, automatic differentiation propagates the directional sensitivity **s**_**v**_(*t*) = *D*_**u**_**x**(*t*; **u**)[**v**] through both the continuous dynamics and the event map. For a bolus expressed in the state units of compartment *k*, with basis vector **e**_*k*_ ∈ ℝ^*n*^ and dose perturbation *v*_*j*_, the jump rule reduces to

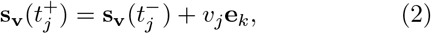

The sensitivity is propagated in the loss function according to

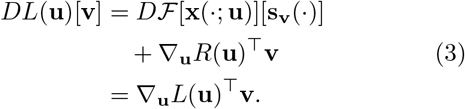

The optimizer therefore receives *L*(**u**) and ∇_**u**_*L* without requiring manually assembled Jacobians, finite-difference approximations, or derivative-free search as the primary optimization engine^27,28,49^.

DiffDose differs from prior dosing optimal control approaches in its control representation and computational interface. Classical biomedical optimal control commonly represents therapy as continuous input and obtains model-specific optimality conditions or adjoint equations^11–14^, whereas DiffDose optimizes an interpretable finite set of administration events. DiffDose differentiates the simulator instead of deriving model-specific sensitivities. OptiDose also implements finite-dimensional dose optimization^24^, but its gradients rely on hand-derived expressions tied to each model and dosing structure. DiffDose retains OptiDose’s interpretable controls while replacing that model-specific derivative engineering with AD through the computational graph.

Differentiating the implemented simulator permits changes to objectives and dosing parameterizations, while preserving gradient-based optimization. Because the simulator remains a mechanistic model rather than an opaque black-box objective, optimized regimens and residual losses can be related back to model parameters, helping to identify patient features associated with dosing decisions and outcomes. To illustrate these advantages, we first evaluated DiffDose against the three OptiDose benchmark studies in Bachmann *et al*.^24^. We then optimized dose timing in a state-dependent DDE, a setting that is inadmissible to OptiDose’s fixed-time dosing representation. Finally, we show that DiffDose’s computational efficiency enables it to scale to individualized QSP virtual population optimization.

In this study, we use DiffDose to denote discretize-then-optimize AD routes (including both forward and reverse modes) through the implemented simulator. To highlight DiffDose’s gradient route advantages, we compared it to continuous sensitivities, adjoints, finite differences while holding the optimizer fixed across gradient routes within each comparison. Downstream optimizers included limited-memory Broyden–Fletcher–Goldfarb–Shanno (L-BFGS), its box-constrained variant (L-BFGS-B)^50^, and adaptive moment estimation (Adam)^51^. Derivative-free comparators included Nelder–Mead^52^ and simulated annealing^53^.

### 2.2 DiffDose forward-mode AD accelerates finite-dimensional dose optimization

To isolate how the differentiation route affects optimization performance, we fixed the numerical simulation, dose schedule, objective, quadrature grid, and optimizer, and varied only the dose-gradient calculation used by DiffDose and its comparators listed above. The benchmarks comprised indirect dose response (IDR), tumor growth inhibition (TGI), and bispecific T-cell engager (BiTE) which were also used by Bachmann *et al*. for OptiDose^24^. Here, we focus on the IDR model as a representative case (see Supplementary Note 1, Supplementary Table 1, and Supplementary Figures 1 and 2 for more details), and provide the details of the other two models in the supplementary material (Supplementary Note 1, Supplementary Tables 2 and 3, and Supplementary Figures 3–6).

In the IDR case, the controlled model output is an exponentially increasing biomarker, *B*(*t*), that is indirectly inhibited (but never extinguished) by a drug (Figure 2a). Six weekly dose amplitudes **u** ∈ [0, 10]^6^ each set seven daily pulses over 42 days. The objective is to keep the biomarker trajectory close to a prescribed target trajectory, *B*_*ref*_ (*t*), minimizing

**Figure 2.**
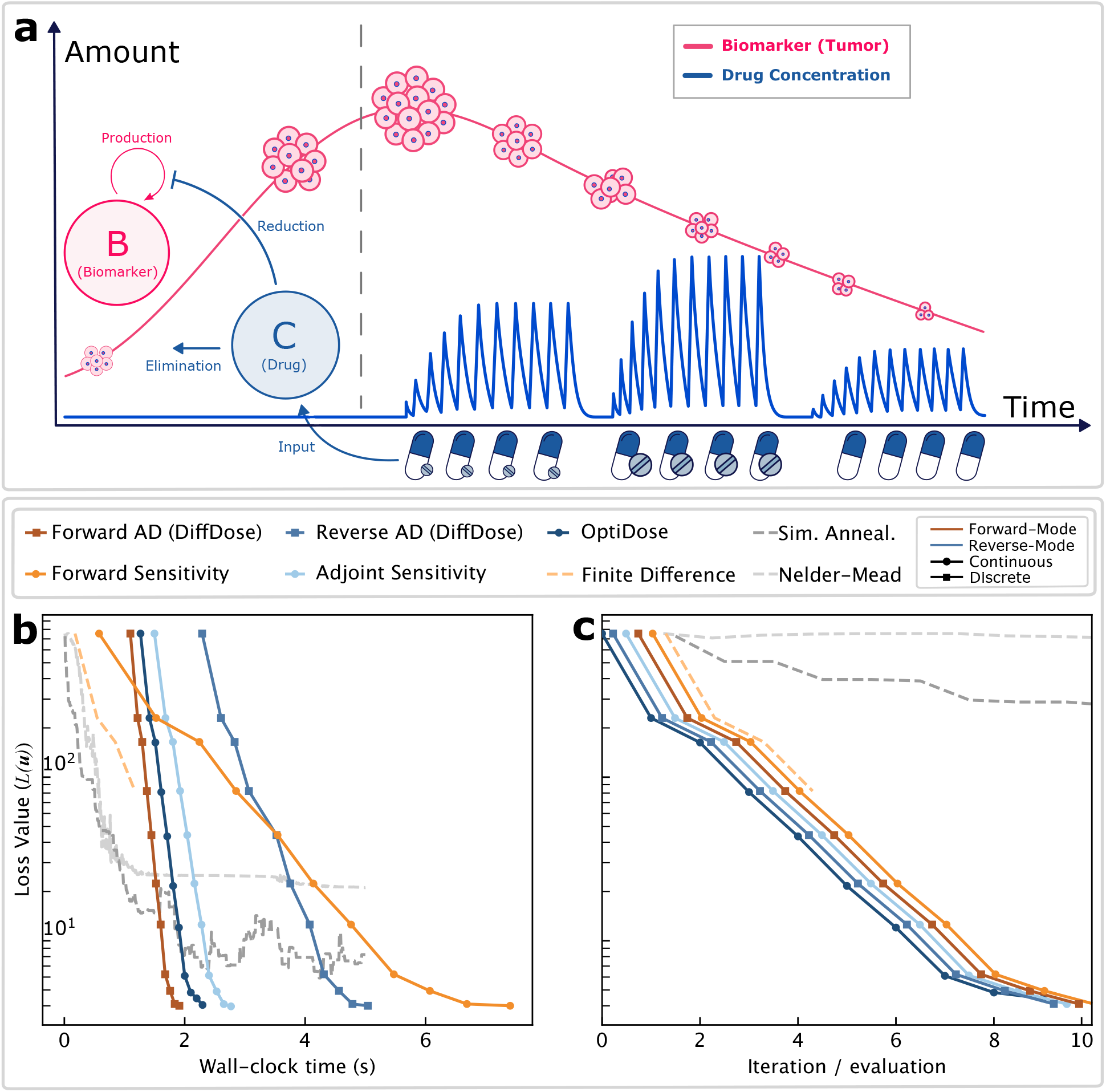
DiffDose forward-mode AD recovers the OptiDose IDR solution with the shortest time-to-solution. **a)** The IDR benchmark uses six dose-amplitude controls at fixed weekly positions; each amplitude sets seven daily pulses that determine drug concentration, C(t), and the indirectly inhibited biomarker, B(t). The vertical axis denotes model-state amount (tumor volume or drug concentration) rather than a shared physical unit. **b–c)** Loss for the same box-constrained problem, **u** ∈ [0, 10]^6^, from **u**_0_ = **1**, shown against (b) wall-clock time and (c) optimizer iteration or objective evaluation. DiffDose-forward AD and DiffDose-reverse AD differentiate through the discretized solver. OptiDose, continuous backsolve adjoint, continuous forward sensitivity, finite differences, and derivative-free searches are comparators. The model, solver, quadrature, objective, and L-BFGS-B configuration were fixed across gradient routes. Most gradient routes reached the same low-loss basin, with DiffDose-forward AD reaching it fastest. Small horizontal offsets in (c) separate otherwise overlapping points.

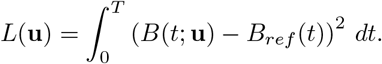

Convergence of DiffDose and the comparators is shown against wall-clock time (Figure 2b) and optimizer iteration or objective evaluation (Figure 2c). The DiffDose forward- and reverse-AD routes, together with the continuous sensitivity and adjoint comparators, reduced the IDR tracking loss from *L*(**u**_0_) ≈ 7.5 × 10^2^ to the order of 10 within about seven quasi-Newton iterations and to the order of 1 within about ten iterations. The continuous backsolve adjoint reached *L* = 3.79; DiffDose forward- and reverse-mode AD agreed to numerical precision at *L* = 4.01; and the OptiDose hand-derived adjoint reached *L* = 4.05. These differences are consistent with each route differentiating a distinct numerical time discretization. Under the same solver settings, finite-difference L-BFGS-B stalled at *L* = 82 after a few iterations because of sensitivity to the finite-difference step size and solver tolerances, while Nelder-Mead plateaued near *L* = 21 and the random-walk Metropolis sampler reached *L* ≈ 5.9.

Runtime separated the gradient routes. DiffDose forward-mode AD reached *L <* 5 in approximately 1.8 s (Figure 2b), whereas DiffDose reverse-mode AD reached the same threshold in approximately 4.6 s, consistent with checkpointing and reverse-mode overhead. The continuous forward-sensitivity comparator recovered the same basin but was slower because it propagated an enlarged system. Direct forward-mode AD therefore best balanced implementation burden and runtime for this six-control problem.

The optimized dose amplitudes were approximately (6.020, 4.057, 0.264, 0.991, 0.773, and 0.860). Because continuous-valued solutions do not directly reflect constraints such as tablet strength, we also implemented a Gumbel–Softmax relaxation over allowable dose levels while retaining end-to-end gradients (Supplementary Note 2). Across IDR, TGI, and BiTE, DiffDose forward-mode AD matched hand-derived adjoint solutions without model-specific Jacobian derivations and gave the shortest time-to-solution for these low-dimensional controls (Supplementary Note 1, Supplementary Tables 1–3, and Supplementary Figures 1–6).

### 2.3 DiffDose optimizes dose-timing in a state-dependent delay model of neutropenia

We next applied DiffDose to optimize granulocyte colony-stimulating factor (G-CSF) timing during cytotoxic chemotherapy, a setting in which fixed-schedule OptiDose is not applicable. Neutrophils are short-lived innate immune cells produced continuously in the bone marrow. Cytotoxic chemotherapy damages rapidly dividing myeloid precursors, lowering absolute neutrophil count (ANC) and increasing infection risk. G-CSF is an endogenous haematopoietic cytokine that promotes neutrophil-lineage differentiation, accelerates maturation, and stimulates release from the marrow reservoir^54,55^ (Figure 3a). We simulated seven CHOP-14 cycles, with cyclophosphamide, doxorubicin, vincristine, and prednisone administered every 14 days^56^, using the state-dependent DDE model of chemotherapy-induced neutropenia developed by Craig *et al*.^46,57,58^. In the CHOP-14 trial^56^, supportive care for neutropenia included exogenous G-CSF; clinical guidelines and protocols recommend short-acting daily filgrastim or its longer-acting pegylated form after chemotherapy^59,60^.

**Figure 3.**
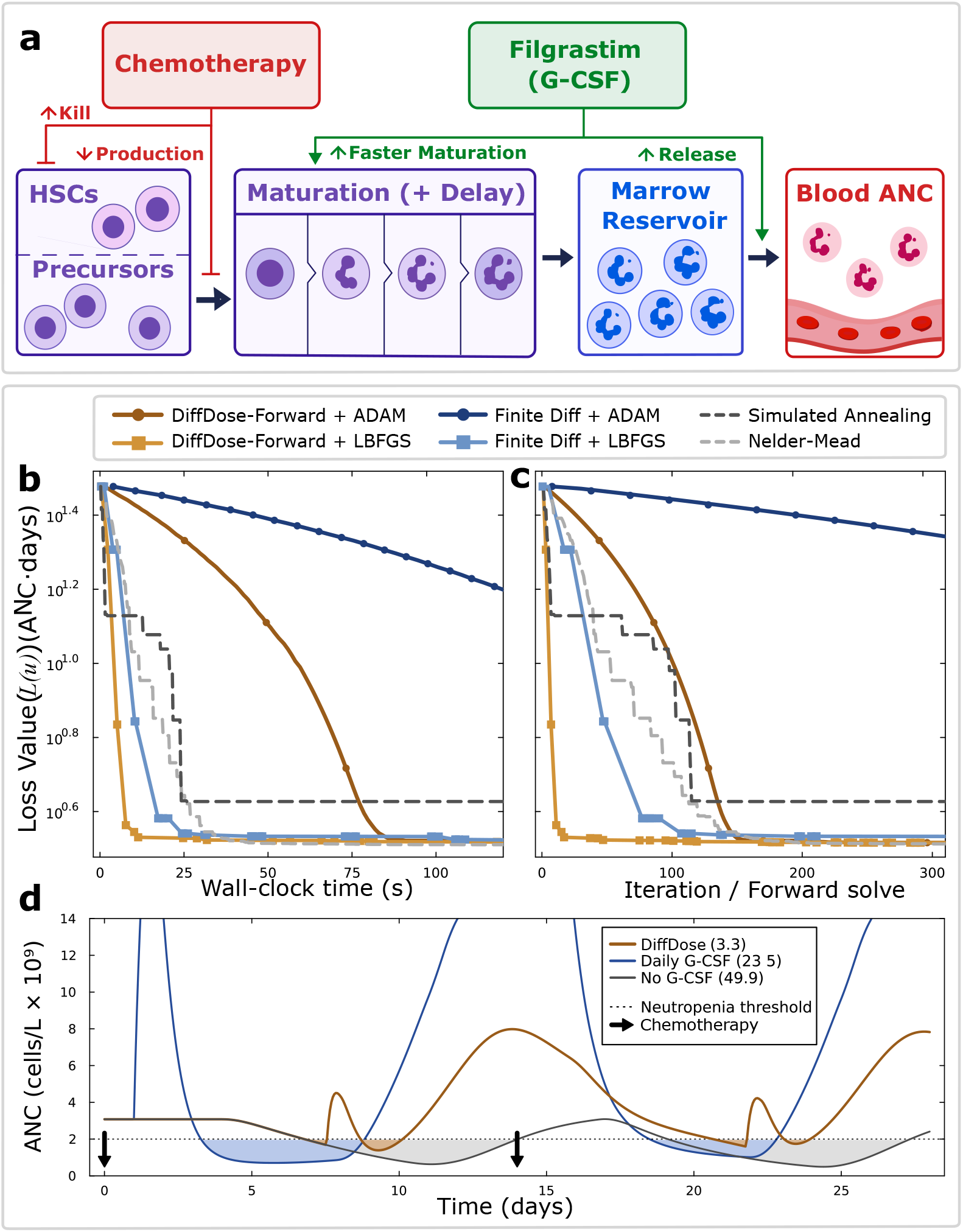
DiffDose identifies a robust day-8 filgrastim window that reduces neutropenia. **a)** Schematic of the state-dependent neutropenia model. Chemotherapy suppresses hematopoietic stem cell and precursor production and increases cell killing, whereas G-CSF accelerates neutrophil progenitor production, neutrophil maturation, and release from the marrow reservoir. **b–c)** Cumulative ANC-deficit loss versus (b) wall-clock time and (c) counted DDE solves when one 300 µg filgrastim administration time per cycle was optimized across seven cycles from the same day-4 initialization. DiffDose-forward AD was paired with Adam or L-BFGS; finite-difference and derivative-free routes served as comparators. **d)** ANC trajectories during the first two cycles under the optimized one-dose-per-cycle schedule, the repeated-daily G-CSF reference, and no G-CSF. Vertical arrows mark chemotherapy administration days, and the horizontal dotted line marks the 2 × 10^9^ cells/L neutropenia threshold. Parenthetical losses cover all seven cycles; the optimized schedule reduced loss by 93.4% versus no G-CSF and 86.1% versus repeated-daily G-CSF.

We asked whether one well-timed 300 µg filgrastim dose per cycle could reduce neutropenia relative to a repeated-daily reference. Here, DiffDose-forward AD and central finite-difference gradients were each paired with Adam or L-BFGS and derivative-free methods served as baselines. DiffDose gradients agreed closely with central finite differences, used as the numerical validation baseline (relative *ℓ*_2_ error 2.9 × 10^−5^). Model provenance, complete equations, gradient validation, and numerical settings are reported in Supplementary Note 3 and Supplementary Tables 4–6; optimization convergence is shown in Figure 3b,c.

DiffDose-forward AD paired with Adam or L-BFGS converged to nearly identical single-dose schedules: approximately 7.4–8.0 days after chemotherapy in each cycle. L-BFGS reached the low-loss basin with fewer optimizer updates and forward solves than Adam (Figure 3b,c), while both reached *L* ≈ 3.28. The best DiffDose-forward AD/L-BFGS schedule had *L* = 3.27, reducing cumulative ANC deficit by 93.4% relative to no G-CSF (*L* = 49.9) and by 86.1% relative to the repeated-daily filgrastim reference^59,60^ on cycle days 1–13 (*L* = 23.5; Figure 3d). A fixed day-8 schedule retained most of this benefit (*L* = 4.387), compared with *L* = 21.04 for random administration days. Within the model, this window is late enough that chemotherapy suppression subsides but precedes prolonged neutropenia, indicating robust timing rather than a need for precise cycle-specific adjustment.

### 2.4 DiffDose optimization in virtual population identifies patient-specific regimens balancing safety and efficacy in bispecific T cell engager therapy

Finally, because individualized dose optimization across a virtual cohort is computationally demanding^61^, we tested whether DiffDose-forward AD could scale to VPops. We used the published 36-state mosunetuzumab QSP model developed by Hosseini *et al*. and Susilo *et al*.^47,48^. Mosunetuzumab is a CD20 (×) CD3 bispecific antibody that brings T cells into contact with malignant B cells. T cell engagers are approved for several hematologic malignancies and selected solid tumors^62^, but can cause IL-6-associated cytokine release syndrome (CRS)^63^. Model and clinical evidence motivated step-up dosing to address the greater cytokine risk during early administrations, when target-cell abundance is high^47^.

For each member of an independently constructed 250-member VPop, DiffDose optimized six dose amplitudes on days 0, 7, 14, 21, 42, and 63 relative to the Hosseini *et al*. regimen. The objective combined day-84 tumor burden, peak IL-6 as a CRS-related safety surrogate, and cycle-level dose regularization (Eq. (7)). DiffDose reduced the objective for every virtual patient, supporting its use within a differentiable in silico trial. Methods, Supplementary Note 4, Supplementary Figures 7–9, Supplementary Tables 7 and 8, and Supplementary Data describe VPop generation, validation, and supporting outcomes.

Relative to the Hosseini *et al*. regimen (i.e., 1.6, 10, 10, 20, 20, and 20 mg), 206 patients (82.7%) improved day-84 residual tumor and 199 (79.9%) reduced the largest IL-6 peak; 163 (65.5%) improved both primary endpoints (Figure 4a,b). Post hoc comparisons showed that 107 patients improved tumor area under the curve (AUC), a measure of cumulative burden, 134 reduced IL-6 AUC, 170 received a lower total dose, and 163 had lower drug exposure. Overall, 156 patients improved at least four of the six reported endpoints, including 23 who improved all six. These counts demonstrate heterogeneous patient-specific trade-offs rather than uniform escalation or dominance.

**Figure 4.**
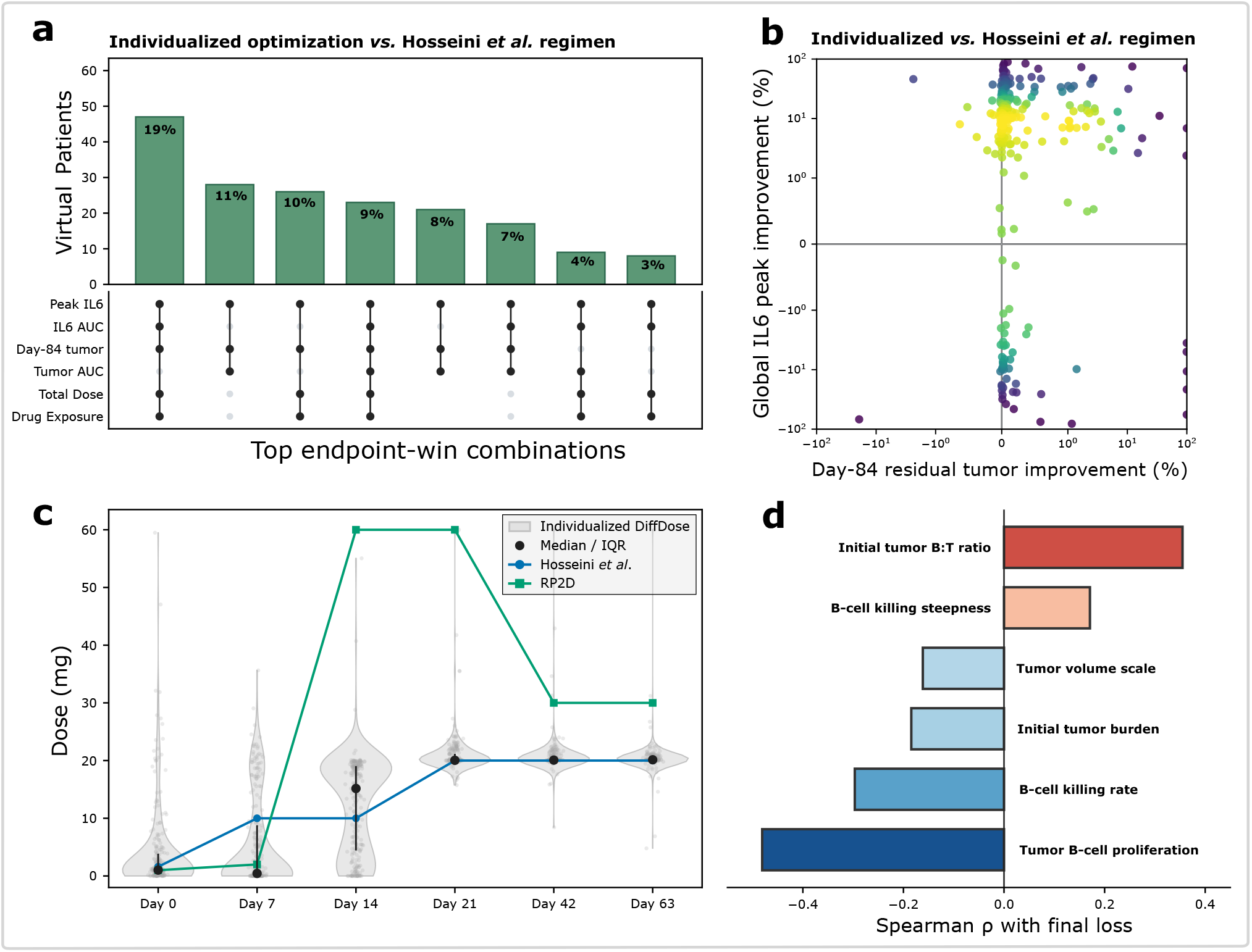
DiffDose identifies heterogeneous, patient-specific mosunetuzumab trade-offs in a virtual population. **a)** UpSet summary of endpoint-win combinations for the 250-member virtual population (VPop) relative to the Hosseini et al. regimen^47^. Bars report the percentage of virtual patients in each combination; linked black dots identify improved endpoints. Lower day-84 tumor burden, tumor AUC, peak IL-6, IL-6 AUC, total dose, and drug exposure were favorable. **b)** Bounded improvements in day-84 residual tumor and largest IL-6 peak, computed for endpoint x as I_x_ = 100(H_x_ − O_x_)/(H_x_ + O_x_), where H and O denote the Hosseini and optimized regimens. Positive values favor DiffDose; medians were 0.34% for tumor and 9.47% for IL-6. Color denotes local point density. **c)** Patient-level optimized doses at the six administration days, with medians, interquartile ranges, and the Hosseini and recommended phase 2 dose schedules. **d)** Spearman rank associations between final DiffDose loss and tumor-related VPop parameters. Positive ρ denotes higher residual loss after optimization; negative ρ denotes lower residual loss. The parameters shown are initial B cell:T cell ratio (`BT_ratio_tumor_init`), B-cell killing response steepness (`nkill`), tumor volume scale (`Vtumor`), initial tumor burden multiplier (`tumor_burden_factor`), B-cell killing rate (`kBkill`), and tumor B-cell proliferation rate (`kBtumorprolif`).

Additional tumor-control gains were often small because 167 patients (67.1%) already achieved at least 99% tumor reduction with the Hosseini regimen, resulting in a median additional day-84 improvement of less than 0.001 percentage points. In contrast, the median IL-6 log ratio was 0.090, corresponding to an 18.8% reduction in the largest IL-6 peak (Figure 4b). This is consistent with the comparison of global IL-6 and tumor-response distributions across fixed and individualized regimens (Supplementary Figure 8) and suggests that simply escalating toward higher-dose schedules is not expected to produce broad additional benefit under the current QSP model, despite the higher target doses in the recommended phase 2 dose (RP2D) schedule near 60 mg per cycle.

The optimized dose distributions provide a mechanistic consistency check (Figure 4c). DiffDose was not constrained to reproduce a step-up schedule, yet it concentrated the widest patient-to-patient variation in cycle 1, kept the first two doses low for many virtual patients, and moved later doses toward the 20 mg target. Variation was greatest at cycle 1 day 15 (C1D15), the point at which the model shifts from early cytokine-risk management toward tumor-control pressure. This pattern is qualitatively consistent with the clinical step-up strategy evaluated by Hosseini *et al*.^47^: early exposure is limited while target-cell abundance and cytokine risk are highest, followed by larger treatment doses. Thus, the gradient-based optimization recovered a clinically interpretable step-up structure from the QSP equations and objective, rather than requiring that structure to be imposed as a fixed candidate regimen.

Parameter associations with final DiffDose loss indicate that residual controllability depends on tumor and immune-response biology (Figure 4d). Virtual patients with faster tumor growth, greater initial burden, or less favorable B cell killing retained higher loss after their six dosing controls were optimized. The strongest associations (most stiff parameter directions) involved tumor B cell proliferation, B cell killing rate and steepness, initial tumor burden, tumor volume scale, and the initial B cell-to-T cell ratio. This pattern is consistent with Susilo *et al*.^48^, who identified tumor size and proliferation, T-cell infiltration and activation, and B cell killing as determinants of mosunetuzumab response. In DiffDose, those features are identified through the local geometry of the optimized dosing problem rather than by testing a small set of prespecified regimens.

## 3 Discussion

DiffDose formulates dose-regimen design as finite-dimensional optimal control over clinically interpretable inputs, and uses differentiable programming to solve these problems in mechanistic pharmacometric and QSP models. Classical workflows choose among continuous adjoints, forward sensitivities, finite differences, and derivative-free search, each with different model and objective requirements. DiffDose, on the other hand, provides a unified interface that supports bolus dosing events, trajectory-dependent and terminal losses, toxicity penalties, dose regularization, and patient-specific targets within one computational graph. This compositional interface of DiffDose is amenable to iterative changes during model development for clinical regimen design, which typically requires balancing efficacy, safety, feasibility, adherence, and operational constraints. Across three complementary settings, DiffDose enabled gradient-based optimization without model-specific hand-derived sensitivities: it matched analytic-adjoint dose-amplitude optima in OptiDose^24^ with improved time to solution, directly optimized timing of G-CSF in a state-dependent DDE model of chemotherapy-induced neutropenia where turnkey adjoints were unavailable, and scaled to a mosunetuzumab QSP VPop, linking residual loss to tumor and immune parameters.

The OptiDose-replication benchmarks showed that DiffDose recovers established dose-control solutions while reducing model-specific derivative work. Conventional workflows require hand-derived adjoints and analytic Jacobians that become restrictive as models, events, and objectives evolve. DiffDose treats the model, events, quadrature, and objective as a composable computational graph, shifting effort from deriving sensitivities to validating the differentiated hybrid dynamical system. In IDR, DiffDose with forward and reverse mode AD reached similar losses. Small differences are expected because continuous-adjoint and discretize-then-optimize gradients can diverge after time discretization, adaptive stepping, interpolation, and event handling^28^. Across IDR, TGI, and BiTE, forward-mode AD gave the shortest time to solution, while AD and adjoint routes reached lower minima than the numerical baselines under matched solver settings.

The neutropenia study extended DiffDose from fixed-time dose amplitudes to administration-time optimization. Timing controls are delicate because moving an event shifts a trajectory discontinuity, requiring explicit event-time derivatives and sensitivity jumps in continuous formulations^64^. Because our dosing-event implementations did not yield validated timing gradients in the state-dependent DDE, we used a smoothed representation that made timing differentiable in the continuous right-hand side. The resulting AD gradients agreed closely with central finite differences, validating the differentiated DDE. Optimized schedules consistently placed G-CSF 7.5–8 days after chemotherapy, consistent with the biological delays in neutrophil production: late enough to avoid stimulating a strongly chemotherapy-suppressed precursor compartment, but early enough to accelerate recovery before prolonged neutropenia. Within the model, a single well-timed injection reduced cumulative ANC deficit more effectively than repeated daily dosing, while a fixed day-8 schedule remained in the optimal basin. Most benefit came from a robust biological window; precise cycle-specific adjustments added little, showing that timing can rival total exposure in nonlinear physiological systems.

At virtual population scale, DiffDose turned the mosunetuzumab QSP model into a differentiable in silico trial. Instead of prespecified fixed regimens, each patient received a personalized six-dose schedule. Benefits were heterogeneous: some patients benefited from lower IL-6 peaks, others from improved tumor control, and some had little room for improvement. This heterogeneity is important because it indicates that the procedure is not merely escalating all patients toward higher exposure. Instead, learned schedules reflect patient-specific tradeoffs encoded by the model and objective. Although no conventional step-up pattern was imposed, early cycle-1 doses remained low for many patients, later doses approached the target dose, and variation peaked at cycle 1 day 15. This pattern is consistent with a transition from early cytokine-risk management, when target abundance and immune activation are high, toward later control of tumor pressure. Recovering this recognizable schedule structure provides a mechanistic consistency check: the QSP dynamics and clinically motivated objective were sufficient to reproduce step-up logic while exposing where dosing was flexible or constrained.

DiffDose also used residual optimized loss to probe controllability. Associations with tumor burden, growth, B-cell:T-cell ratio, and B-cell killing identified virtual patients that remained difficult to control after all six dosing degrees of freedom were used. The optimized objective therefore measures residual treatment difficulty. This differs from sensitivity analysis under a fixed regimen: a plausible parameterization can occupy a different local dynamical regime once its patient-specific dose vector is optimized. Incorporating dose optimization into virtual population evaluation and generation may distinguish descriptive variability from variability that directly constrains treatment controllability. More broadly, differentiable in silico trials can answer counterfactual questions about both regimen and cohort composition.

Clinical trials are themselves empirical methods for dose regimen optimization, while off-label uses of approved drugs personalize those same medicines. DiffDose preserves an end-to-end mechanistic link between dose decisions and patient features and exposes the local geometry of the patient-parameter manifold around each optimal controller. Evaluating model robustness in this neighborhood may inform cohort selection, response stratification, and prospective dosing hypotheses.

Several limitations temper these findings. Every optimized regimen depends on the correctness of the underlying mechanistic model and its implementation in an AD-compatible numerical stack; legacy workflows, including many MATLAB models, may require refactoring into differentiable JAX or Julia solvers^65^. Objective targets, weights, and constraints determine the selected efficacy, safety, and dose trade-offs; a different clinical objective can select a different regimen even with the same model and gradient route. These are also non-convex local problems. L-BFGS and local optimizers exploit accurate gradients but do not guarantee a global optimum, so multi-start or global-local methods may be needed when several clinically distinct basins exist. Gradients through adaptive solvers depend on tolerances, interpolation, event handling, and smoothing, making validation against independent numerical routes essential. These differences reflect the discretized numerical program that AD differentiates.

Future work should extend DiffDose in three directions. First, residual, patient-parameter, model-form, and VPop uncertainty should enter robust or Bayesian objectives. Second, closed-loop dosing could update controls as biomarkers, toxicity, imaging, or laboratory measurements arrive, including through nonlinear model-predictive control (NMPC). Third, differentiable simulators could train neural network parameterized policies end-to-end rather than optimizing each regimen independently. These learned functions can map patient states, baseline covariates, or inferred latent parameters to dose decisions, while the mechanistic QSP or PK/PD model supplies the differentiable training environment. Characterizing the local geometry of the joint patient-parameter and controller landscape near these optima may reveal sloppy or stiff VPop directions. These extensions require explicit safety constraints, uncertainty calibration, and prospective validation, and could connect mechanistic pharmacology with control and data-adaptive precision dosing.

In summary, our results support DiffDose as a differentiable programming solution for mechanistic dose-regimen optimization. Our approach reproduced established ODE dose-amplitude benchmarks, enabled timing optimization in a state-dependent DDE, and scaled to individualized optimization in a QSP VPop. It makes mechanistic models actionable as optimization engines. By differentiating dosing variables, hybrid dynamics, and clinically aligned objectives together, DiffDose provides a reusable basis for individualized regimen design and more informative in silico clinical trials.

## 4 Methods

### 4.1 Numerical simulation and differentiation approaches

Each objective evaluation involved two common operations: solve the mechanistic model for a proposed finite dose vector, then compute trajectory-derived endpoints and a scalar loss. Gradient-based routes additionally differentiated that loss with respect to the controls. The ODE and DDE models were solved by adaptive time integration. Fixed-time doses were represented as state-reset maps that instantaneously add dose to a designated compartment, except for the matched short-window OptiDose input; the DDE timing study used a smooth filgrastim gate, so administration time entered the continuous right-hand side. For each head-to-head gradient comparison, the model, dosing representation, solver, loss, bounds, initialization, and downstream optimizer were fixed. Case-specific algorithms, tolerances, output grids, and validation tests are given below and in Supplementary Notes 1, 3 and 4.

We compared two differentiation viewpoints^27,28,49^. DiffDose uses discretize-then-optimize AD: forward mode propagates Jacobian-vector products (JVPs), and reverse mode propagates vector-Jacobian products (VJPs), through the implemented solver and loss. Continuous sensitivity and adjoint equations are optimize-then-discretize comparators; finite differences and derivative-free methods are numerical baselines. Within each benchmark, gradient routes supplied the same bounded optimizer, and no dense Jacobian was assembled for the DiffDose routes. The route taxonomy, equivalent variational, Lagrange-multiplier, and Pontryagin maximum-principle interpretations, and optimizer interfaces are summarized in Supplementary Note 5.

For fixed-time ODE controls, DiffDose differentiates the hybrid simulation defined in Eq. (1), including the state-reset map at each administration. For an additive bolus, differentiating the reset gives the jump rule in Eq. (2). AD evaluates Jacobian-vector or vector-Jacobian products (JVPs or VJPs). For example, *D*_**x**_*f* (*t*, **x, p**) **s** is the action of the local linearization on a tangent vector **s**, without a preassembled symbolic or dense Jacobian matrix^27,28^. The resulting objective-gradient pair is passed to the optimizer. Event-sensitive derivations and their relationship to continuous sensitivity and adjoint formulations are retained below for completeness and expanded in Supplementary Note 5.

Within DiffDose, gradients of the objective, *L*(**u**), are propagated through the fixed-time hybrid ODE. Let **v** ∈ ℝ^*m*^ be an arbitrary perturbation of the dose vector, **u**. The corresponding directional state sensitivity is defined as

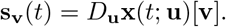

This tangent variable gives the first-order perturbation of the hybrid trajectory induced by **v**, the dose perturbation^66^. Since the vector field is allowed to be arbitrarily nonlinear in **x**, the tangent equation is obtained by linearizing the nonlinear dynamics along the current trajectory

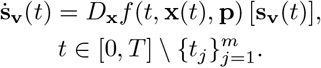

At a dosing event, the directional sensitivity

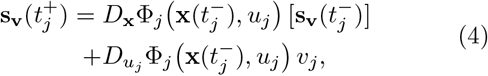

is propagated by differentiating the event function^29^. In DiffDose, we considered dosing events that directly increment the value of a state variable at time *t*_*j*_ by adding a bolus dose amount 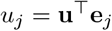, where **e**_*j*_ ∈ ℝ^*m*^ is the *j*-th Euclidean basis vector in control space; *k* indexes the dosing compartment of the ODE system. We set the dose administration event function to be Φ_*j*_(**x, u**) = **x** + **u**^⊤^**e**_*j*_ **e**_*k*_, with **e**_*k*_ ∈ ℝ^*n*^. Thus, we have the identity Jacobian with respect to **x** ∈ ℝ^*n*^, *D*_**x**_Φ = **I**_*n*_ ∈ ℝ^*n*×*n*^ and the basis Jacobian with respect to the control variable at the control time, 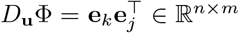. The resulting Jacobian actions are therefore

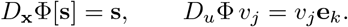

For this additive reset, the general event rule in Eq. (4) reduces to Eq. (2); other reset maps use their corresponding derivatives. Including the direct dose penalty, the objective derivative is given by Eq. (3).

In practice, the components of ∇_**u**_*L* may be obtained by seeding the tangent equation with the coordinate perturbations **v** = **e**_*j*_ or by propagating a batch of such seed directions through the automatic differentiation backend (i.e., dual-number chunking). This is equivalent to using the full sensitivity matrix **S**(*t*) = *D*_**u**_**x**(*t*; **u**), but the calculation does not require analytic or numerically approximated Jacobian matrices. The resulting pair (*L*(**u**), ∇_**u**_*L*(**u**)) is instead passed to a numerical optimizer (e.g., quasi-Newton method).

The above approach is termed discretize-then-optimize, since we compute the objects needed for the gradient at each discretization of the solution of the ODE problem. We refer the reader to Supplementary Note 5 for the optimize-then-discretize version of the gradient derivation. Note that the adjoint dual of this sensitivity formulation is continuous without any jumps at additive dose events and the gradient of the loss function with respect to the dose amplitude is a lookup of the adjoint at the right limit of the dose time (*t*^+^).

### 4.2 Benchmarking DiffDose for dose-amplitude control

Bachmann *et al*. developed OptiDose to optimize discrete-dose controllers for ODE models of pharmacological systems. Their optimize-then-discretize scheme derives dose gradients from Karush–Kuhn–Tucker conditions and uses analytic Jacobians assembled manually for each ODE model^31,44^. OptiDose remains one of the few implementations of finite-dimensional optimal control of administered doses for pharmacology^24^. To benchmark DiffDose against this reference, we reproduced the three PK/PD problems reported by Bachmann *et al*.^24^: the indirect biomarker response (IDR) model, tumor growth inhibition (TGI) model, and bispecific T-cell engager (BiTE) model. OptiDose represents each administration as a smooth Dirac-impulse approximation over a short *ε*-window; DiffDose additionally evaluates direct AD through the discretized simulation.

In our notation, for one control per administration, the dose gradients reduce to integrals of the adjoint component within the *ε*-window at the injection coordinate:

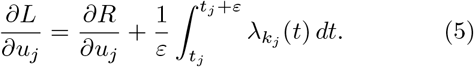

Here, *λ*_*k*_ is the adjoint component for the dosing compartment, defined in Supplementary Note 1. Shared-amplitude expressions are given in Supplementary Note 1. We used the corresponding model-specific expressions as the analytic-adjoint references. The OptiDose reference and DiffDose differentiation routes were implemented with matched settings in JAX/Diffrax^67^. We compared routes using cumulative wall-clock time, with all six-control IDR runs beginning from **u**_0_ = (1, …, 1) with bounds [0, 10]^6^, and each gradient route supplied to the same L-BFGS-B configuration. Timings included solver, differentiation, line-search, and optimizer work after one untimed JAX tracing/JIT call. This end-to-end timing therefore captures the numerical integrations required by each continuous or discrete differentiation route

Complete equations, parameters, reference trajectories, analytic adjoints, and additional IDR, TGI, and BiTE results are provided in Supplementary Note 1 and Supplementary Tables 1–3.

### 4.3 DiffDose timing optimization in a state-dependent neutropenia DDE

The DDE timing study comprised three implementation steps: translating the published state-dependent model, introducing a differentiable administration-time parameterization, and comparing matched downstream optimizers.

We translated the published state-dependent DDE model of granulopoiesis and chemotherapy-induced neutropenia developed by Craig *et al*. from MATLAB to Julia^46,57,58^. The model comprises four coupled modules: (1) delayed hematopoietic stem-cell renewal and differentiation; (2) delayed neutrophil precursor amplification, maturation, and reservoir release; (3) two-compartment G-CSF PK/PD; and (4) four-compartment chemotherapy PK/PD. Chemotherapy was represented as an intravenous infusion that suppresses self-renewing hematopoietic stem cells and neutrophil precursors, whereas filgrastim was represented as subcutaneous exogenous G-CSF^54,55^. Complete equations, model states, and the initial history are given in Supplementary Note 3 and Supplementary Tables 4 and 5. In the source model, parameter values were both fixed to published values and estimated during the original calibration; quantities constrained by steady-state conditions are recalculated at homeostasis by the Julia constructor. Exact values, defining calculations, and implementation provenance are available in the DiffDose repository described in the data and code availability statements. Treatment and numerical settings are reported in Supplementary Table 6.

Here, the clinical objective is to reduce neutropenia. Hence, we defined the loss function to account for neutrophil deficit as the cumulative shortfall of ANC below the neutropenia threshold *N*_thr_ = 2 × 10^9^ cells/L^68^:

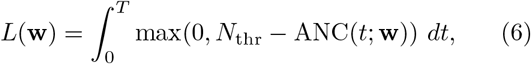

where ANC(*t*) is the absolute neutrophil count at time *t* and the decision vector, **w**, denotes the time post-chemotherapy (per cycle) at which exogenous G-CSF treatment begins.

The DelayDiffEq.jl implementation^69^ was differentiated directly with ForwardDiff.jl^70^. In the software versions used here, SciMLSensitivity.jl^71^ did not provide turnkey continuous forward-sensitivity or adjoint equations for this state-dependent DDE. Continuous variational and adjoint formulations exist for state-dependent DDEs but require model-specific differentiation of the lag^72^. We therefore used DiffDose forward AD on the smoothed, discretized objective and validated its gradients against central finite differences. ForwardDiff gradients and finite differences were each supplied to Adam or L-BFGS; Nelder–Mead and simulated annealing were derivative-free comparators. Adam used projected updates, whereas L-BFGS used Optim.jl Fminbox^73^. All methods began from **w**_0_ = (4, …, 4) days to match the prior model-analysis setup. Supplementary Note 3 details the Julia implementation of the published model, the smooth timing parameterization, and the complete model and solver settings.

### 4.4 DiffDose optimization in a mosunetuzumab QSP virtual population

To implement the virtual population study, we translated the published 36-state mosunetuzumab ODE model of Hosseini *et al*. and used the virtual-patient analysis of Susilo *et al*. to guide biological variability^47,48^. Fixed-time doses were represented as instantaneous state resets, and outputs retained for optimization included tumor burden, IL-6, drug exposure, and B and T cell states and activation. Because the original VPop parameters were unavailable, we screened 62 inputs with the extended Fourier amplitude sensitivity test (eFAST)^74^, retained stable and biologically curated directions, resimulated 20,000 candidates, and selected 250 parameterizations by stochastic pruning. The selected cohort matched digitized IL-6, activated T-cell, day-84 tumor, and responder-fraction summaries reported by Hosseini *et al*. while preserving parameter diversity. Supplementary Note 4 and Supplementary Figure 7 describe cohort construction and validation. Supplementary Data gives the exact 250-member VPop parameterizations; the complete sampling specification and bounds are available in the DiffDose repository.

To apply DiffDose to the QSP setting, we optimized six independent dose amplitudes for each virtual patient at fixed administration days:

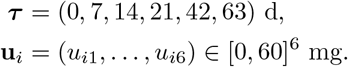

Six dose amplitudes were optimized over four 21-day cycles: three step-up doses on days 1, 8, and 15 of cycle 1 (C1D1, C1D8, and C1D15), followed by one dose on day 1 of cycles 2–4 (C2D1–C4D1). Taking C1D1 as simulation day 0, the fixed administration times were ***τ*** = (0, 7, 14, 21, 42, 63) days.

Rather than optimizing against a single fixed reference regimen, we defined patient-specific achievable efficacy and safety targets. For each virtual patient, the tumor control target was taken to be the ratio of the predicted day-84 tumor size to initial tumor size (i.e., day-84 tumor ratio) obtained under fixed maximal dosing, whereas the cytokine target was the IL-6 peak obtained under minimal dose amount, with IL-6 used as a continuous pharmacodynamic safety surrogate, not as a clinical CRS diagnostic threshold. We therefore optimized the loss function:

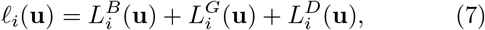

where 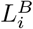 is the one-sided day-84 tumor endpoint penalty, 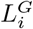 is the one-sided global IL-6 peak penalty, and 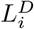 regularizes the cycle-level dose totals toward 20 mg. Hence, the individualized objective penalizes (1) failure to elicit a patient’s maximal possible tumor response, (2) failure to preserve the patient’s low-dose IL-6 safety profile, and (3) deviations from a regimen of approximately 20 mg per cycle, as in Hosseini *et al*. Full definitions of the endpoint targets, normalization constants, and smooth one-sided penalties are given in Supplementary Note 4. Then, for each virtual patient, we solved

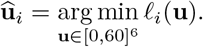

For each of the 250 virtual patients, the 84-day ODE was solved with Rodas4P (absolute tolerance 10^−8^; relative tolerance 10^−5^) and differentiated with ForwardDiff.jl through the solver and endpoint calculation. The six doses were optimized with Fminbox(BFGS()) from an independent Uniform(0, 60)^6^ initialization. We retained the lowest loss finite dose vector in the logged history. We also evaluated alternative objective families, including Hosseini-reference and RP2D-reference step-up losses as well as a cohort-shared problem in which one common six-dose regimen minimizes mean loss across the VPop, as formalized in Supplementary Note 4 and Supplementary Table 8.

## Code availability

The OptiDose benchmarks were implemented in JAX and Diffrax^67^; the neutropenia DDE and mosunetuzumab QSP studies used Julia with DelayDiffEq/SciMLSensitivity, ForwardDiff, and Optim.jl^69–71,73^. Gradient construction was kept separate from each downstream optimizer. Supplementary Note 5 summarizes the differentiation and optimizer interfaces, while Supplementary Notes 1, 3 and 4 give case-specific solver settings. Code used to reproduce the PK/PD benchmarks, neutropenia DDE, mosunetuzumab virtual population optimization, and figures is available under the BSD 3-Clause License at https://github.com/Craig-Lab/DiffDose.

## Data availability

Supplementary Data contains the parameter vectors for all 250 virtual patients used in the mosunetuzumab analyses. The complete 62-parameter sampling specification, including bounds and sampling scales, is available with the analysis code at https://github.com/Craig-Lab/DiffDose.

## Author contributions

SH, MC, and AE conceived the project. MC and AE supervised the project and obtained funding. SH conducted ideation, design, implementation, optimization, and validation. SH, MC, and AE wrote the first draft of the manuscript. All authors revised and approved the final manuscript.

## Acknowledgements

This work was supported by the New Frontiers in Research Fund (NFRFE-2021-00336) and NSERC (RGPIN2018-04546 to MC and RGPIN-2019-04460 to AE). SH was funded by an Alexander Graham Bell Canada Graduate Scholarship–Doctoral (CGS-D-579679-2023). MC is the Canada Research Chair in Computational Immunology, and this research was undertaken, in part, thanks to funding from the Canada Research Chairs Program.

## Supplementary Information

### Guide to the Supplementary Information

This PDF contains five Supplementary Notes with supporting results, model definitions, numerical methods, validation analyses, and implementation details for the three case studies in the main manuscript. Supplementary Data provides the selected virtual population parameter matrix. The complete virtual population sampling specification and bounds, neutropenia delay differential equation (DDE) parameterization and implementation, translated mosunetuzumab quantitative systems pharmacology (QSP) model, and associated analysis code are versioned in the public DiffDose repository: https://github.com/Craig-Lab/DiffDose.

**Supplementary Note 1:** ODE benchmark results and implementation

**Supplementary Note 2:** Differentiable relaxation of quantized dose controls

**Supplementary Note 3:** Neutropenia DDE validation and implementation

**Supplementary Note 4:** Mosunetuzumab virtual population construction and optimization

**Supplementary Note 5:** Differentiation routes and downstream optimization

### Supplementary Note 1: ODE benchmark results and implementation

#### Indirect biomarker response benchmark

The indirect biomarker response (IDR) benchmark compared DiffDose forward- and reverse-mode automatic differentiation (AD) with the analytic OptiDose adjoint, a continuous backsolve adjoint, an explicit forward-sensitivity system, finite differences, and derivative-free baselines. All gradient routes used the limited-memory Broyden–Fletcher–Goldfarb–Shanno algorithm with box constraints (L-BFGS-B), with the same model, objective, bounds, initialization, solver, quadrature, and optimizer settings. Supplementary Table 1 reports the terminal benchmark values; Supplementary Figures 1 and 2 show the optimized trajectories and the comparison between smooth and jump representations, respectively.

**Supplementary Table 1.**
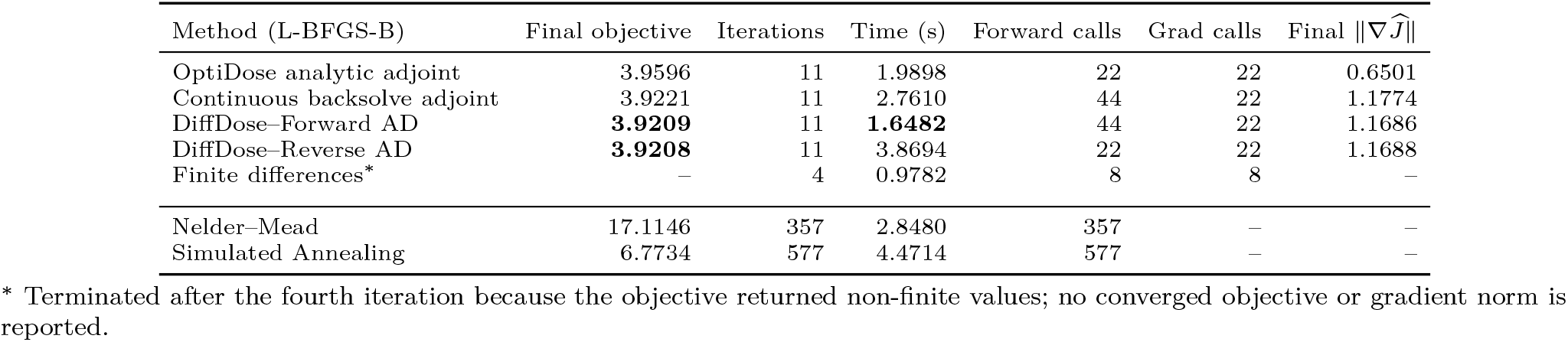
Gradient-route and baseline comparisons for the IDR benchmark. Bold values mark the lowest final objective and shortest wall-clock time among converged gradient-based routes, with ties based on the displayed precision. L-BFGS-B denotes the limited-memory Broyden–Fletcher–Goldfarb–Shanno algorithm with box constraints. The horizontal rule separates derivative-free methods, which do not use gradients.

**Supplementary Figure 1.**
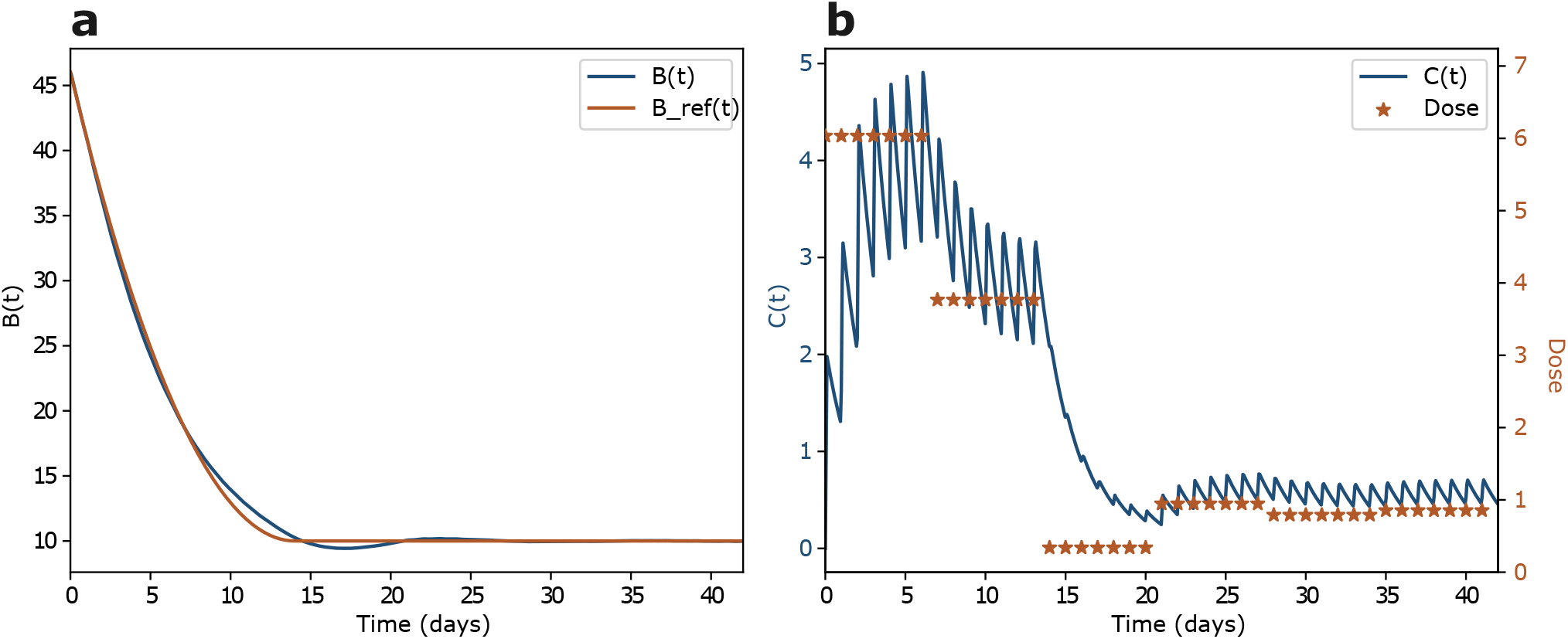
Optimized IDR trajectories obtained with DiffDose–Forward AD. (a) The biomarker trajectory B(t) follows its prescribed reference B_ref_(t). (b) Drug concentration C(t) under the six optimized dose-amplitude controls. Dose markers indicate the fixed administration times.

**Supplementary Figure 2.**
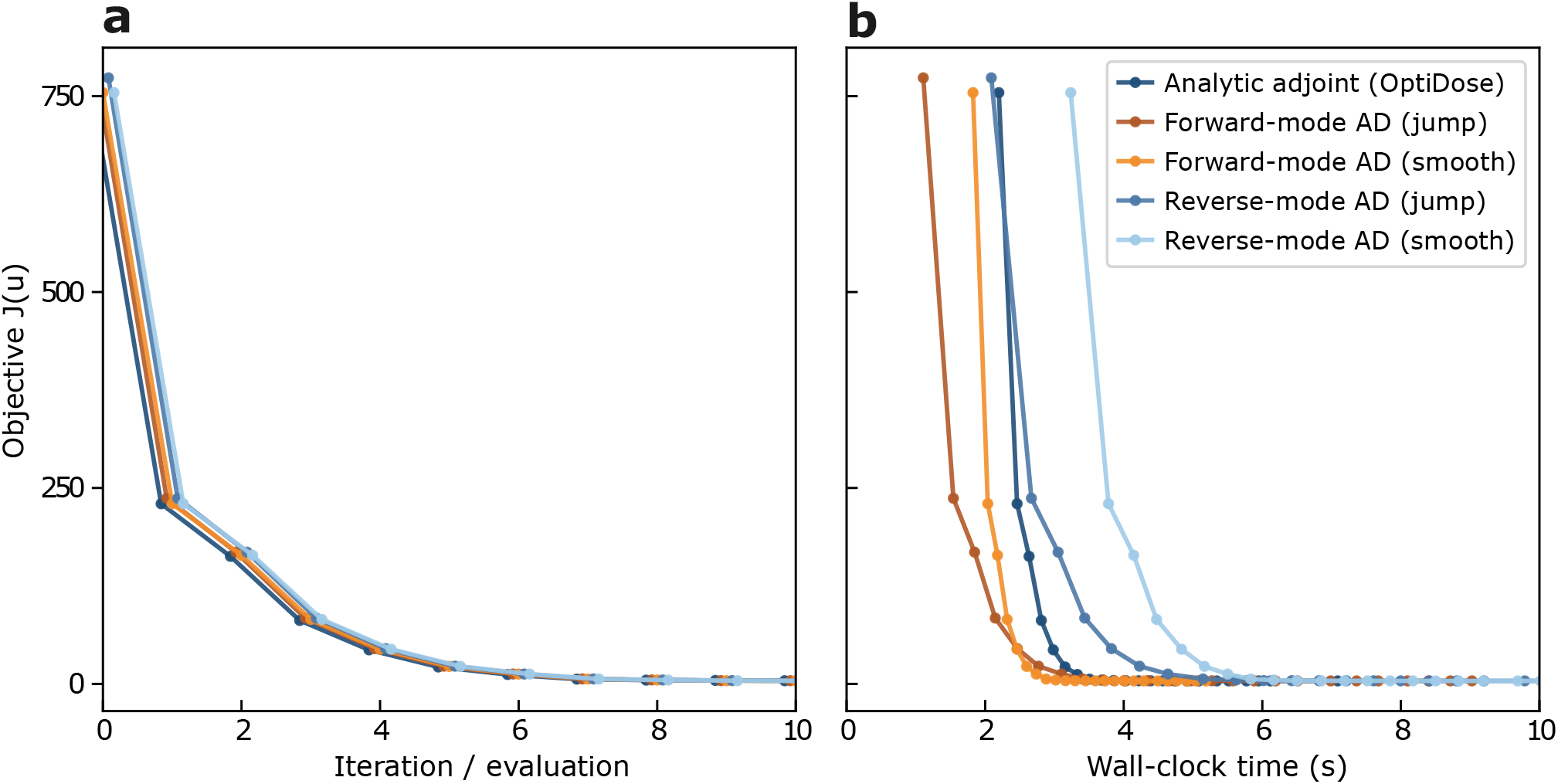
Smooth and jump dosing representations recover similar IDR solutions. (a) Objective J (**u**) versus optimizer iteration or evaluation. Markers are shifted slightly along the horizontal axis to show overlapping values. (b) Objective versus wall-clock time. Smooth routes represent each bolus by a short infusion window; jump routes apply instantaneous state updates at the same fixed administration times. All routes used the same initial dose vector and bounds, and jump-route solutions were re-evaluated with the common smooth objective.

#### Tumor growth inhibition benchmark

The tumor growth inhibition (TGI) benchmark tested the same gradient routes in a larger pharmacokinetic/pharmacodynamic (PK/PD) model. Supplementary Table 2 summarizes the comparison, and Supplementary Figures 3 and 4 show convergence and the corresponding optimized trajectories.

**Supplementary Table 2.** Gradient-route and baseline comparisons for the TGI benchmark. Bold values mark the lowest final objective and shortest wall-clock time among converged gradient-based routes. L-BFGS-B denotes the limited-memory Broyden–Fletcher–Goldfarb–Shanno algorithm with box constraints. The horizontal rule separates derivative-free methods, which do not use gradients.

| Method (L-BFGS-B) | Final objective | Iterations | Time (s) | Forward calls | Grad calls | Final $\ \nabla \hat{J}\ $ |
| --- | --- | --- | --- | --- | --- | --- |
| OptiDose analytic adjoint | **0.0028129** | 28 | 4.7034 | 28 | 28 | – |
| Continuous backsolve adjoint | 0.0028138 | 28 | 6.1210 | 28 | 28 | – |
| DiffDose–Forward AD | 0.0028139 | 28 | **4.2597** | 28 | 28 | – |
| DiffDose–Reverse AD | 0.0028139 | 28 | 7.8476 | 28 | 28 | – |
| Finite differences | 0.0028139 | 28 | 15.2259 | 28 | 28 | – |
| Nelder–Mead | 0.0258805 | 940 | 15.0161 | 940 | – | – |
| Simulated Annealing | 0.0531493 | 739 | 15.0050 | 739 | – | – |

**Supplementary Figure 3.**
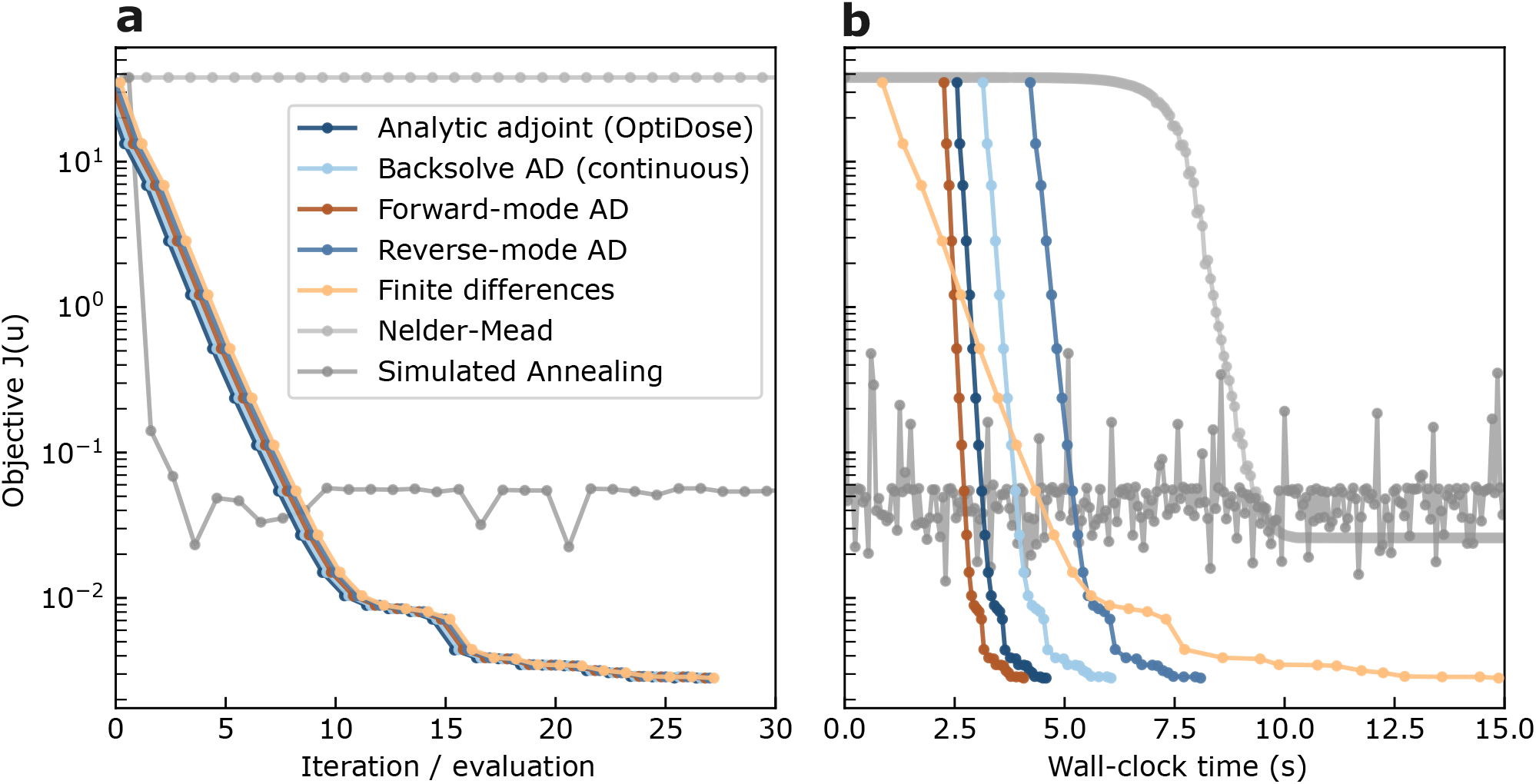
DiffDose reaches the OptiDose TGI solution with lower computational cost. (a) Log-scaled objective J (**u**) versus optimizer iteration or evaluation. Markers are shifted slightly along the horizontal axis to show overlapping values. (b) Objective versus wall-clock time. All gradient routes used the same model, objective, dose bounds, initialization, solver settings, and L-BFGS-B configuration; derivative-free methods are shown as baselines.

**Supplementary Figure 4.**
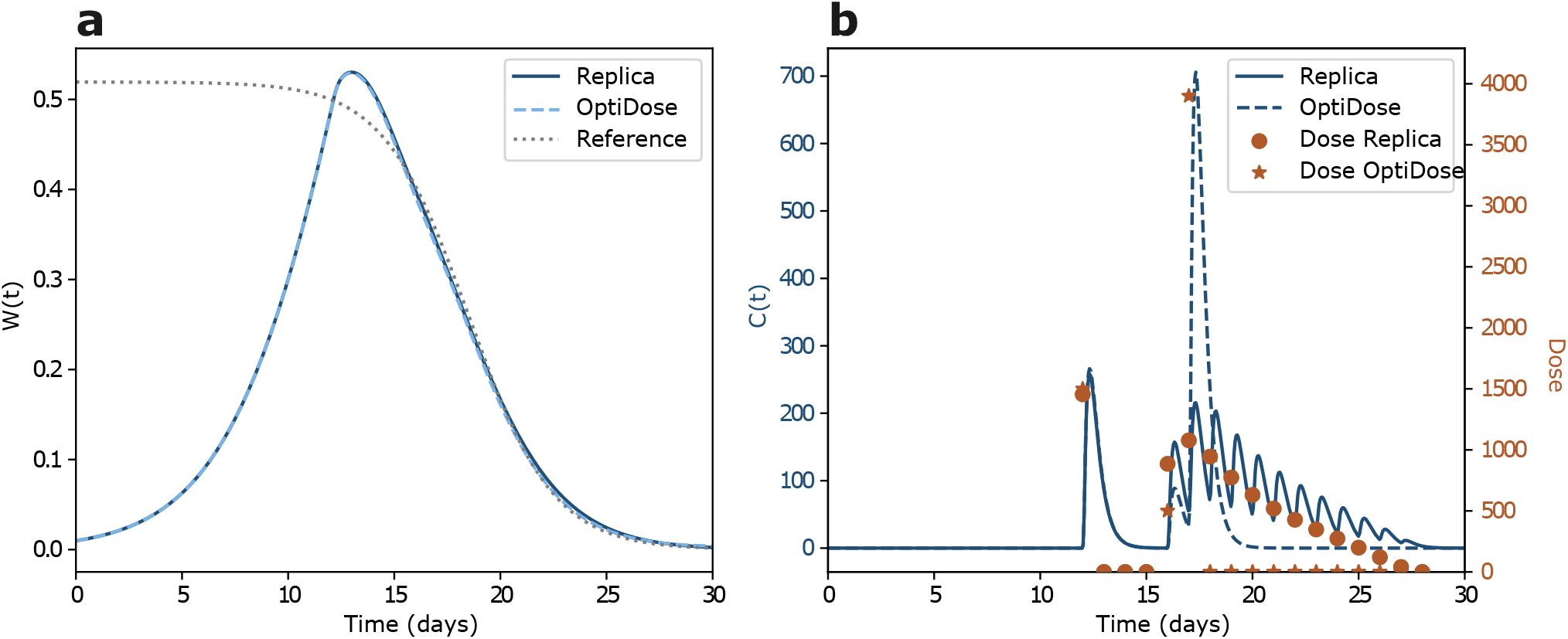
Optimized TGI trajectories. (a) Tumor-size trajectories obtained with the replicated OptiDose and DiffDose solutions relative to the target trajectory. (b) Drug concentration and fixed-time dose amplitudes for the two optimized solutions.

#### Bispecific T cell engager benchmark

The bispecific T cell engager (BiTE) benchmark tested a single repeated dose-amplitude control in a target-mediated model. Supplementary Table 3 reports the optimization results, and Supplementary Figures 5 and 6 show convergence and the optimized state trajectories.

**Supplementary Table 3.** Gradient-route and baseline comparisons for the BiTE benchmark. Bold values mark the lowest final objective and shortest wall-clock time among converged gradient-based routes. L-BFGS-B denotes the limited-memory Broyden–Fletcher–Goldfarb–Shanno algorithm with box constraints. The horizontal rule separates derivative-free methods, which do not use gradients.

| Method (L-BFGS-B) | Final objective | Iterations | Time (s) | Final $u$ | Final $\ \nabla \hat{J}\ $ |
| --- | --- | --- | --- | --- | --- |
| OptiDose analytic adjoint | **5.256155** | 11 | 170.0287 | 663.8549 | $6.95 \times 10^{-9}$ |
| DiffDose–Forward AD | 5.256156 | 10 | **37.3733** | 663.7893 | $5.31 \times 10^{-9}$ |
| DiffDose–Reverse AD | **5.256155** | 11 | 100.6156 | 663.8549 | $6.95 \times 10^{-9}$ |
| Finite differences* | 5.256249 | 8 | 70.3734 | 662.8107 | $2.98 \times 10^{-5}$ |
| Nelder–Mead | 5.256156 | 50 | 46.9577 | 663.8477 | – |
| Simulated Annealing | 5.508096 | 14 | 15.9973 | 713.7436 | – |

**Supplementary Figure 5.**
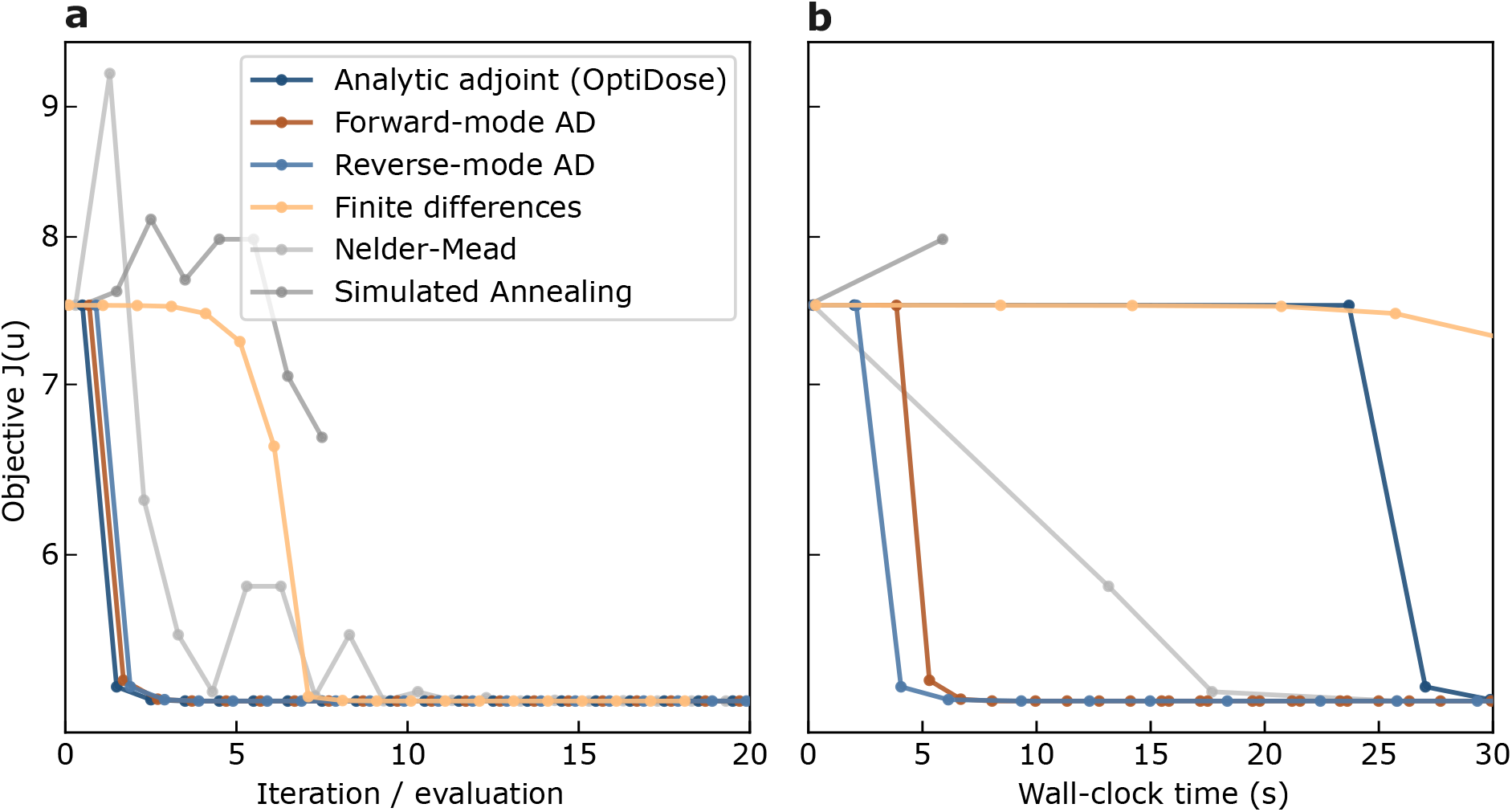
DiffDose recovers the OptiDose solution in the BiTE benchmark. (a) Log-scaled objective, J (u), versus optimizer iteration or evaluation. Markers are shifted slightly along the horizontal axis to show overlapping values. (b) Objective versus wall-clock time. All methods used the same model, objective, dose bounds, initialization, solver settings, and downstream optimizer where applicable.

**Supplementary Figure 6.**
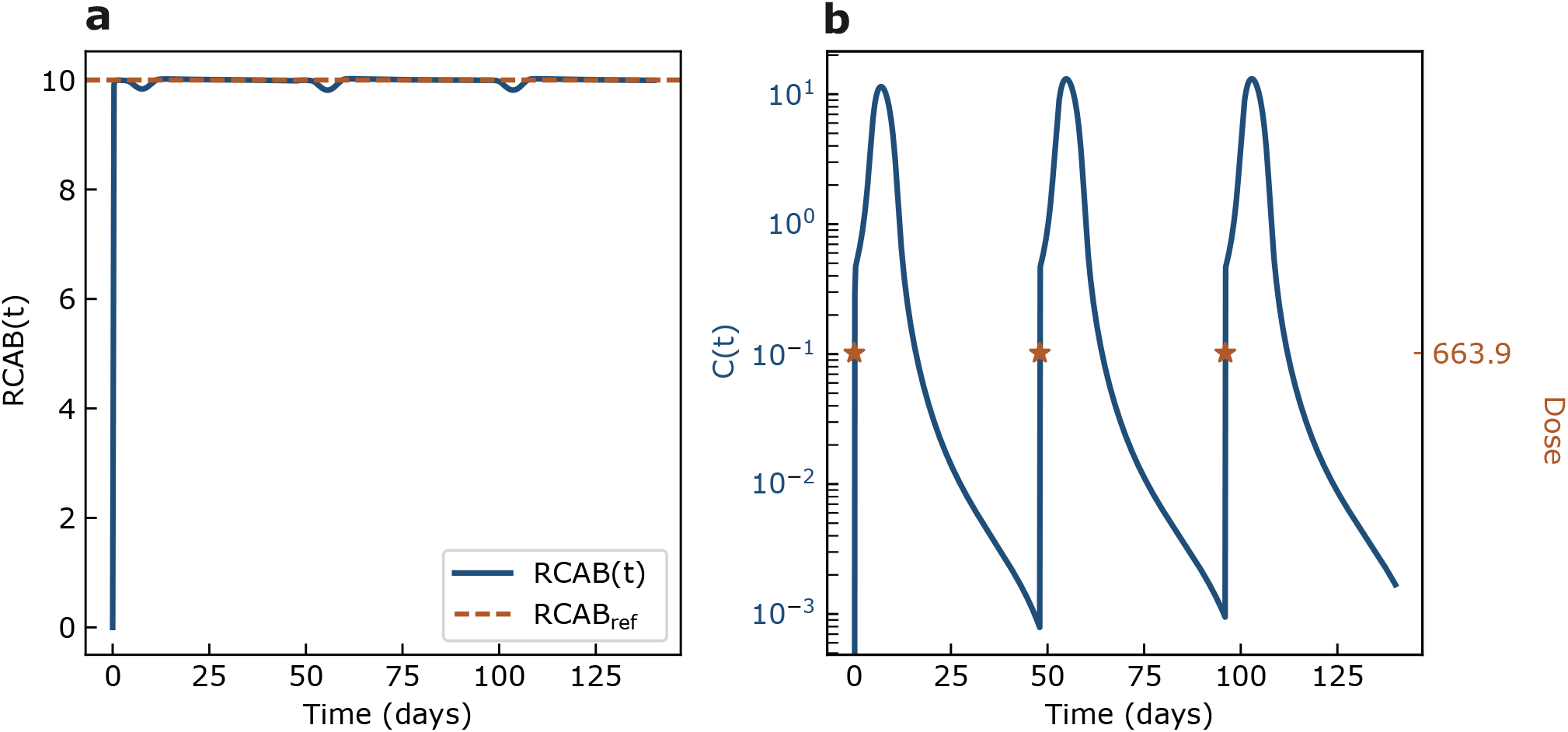
BiTE state trajectories at the optimized dose. (a) The controlled RC_AB_ (t) state relative to its reference R_ref_. (b) Drug concentration at the repeated optimized dose of 663.9, obtained with the OptiDose analytic-adjoint solution and numerically matched by DiffDose.

#### Benchmark formulation and model definitions

The IDR, TGI, and BiTE equations, parameter values, dosing schedules, and target trajectories below reproduce the published OptiDose benchmarks of Bachmann *et al*.^1^ Our changes concern their implementation in JAX/Diffrax and the routes used to compute dose gradients; the benchmark definitions were otherwise retained unless stated explicitly.

##### General finite-dimensional optimal dosing problem

For a given dosing strategy *u*, the state trajectory is determined by the ordinary differential equation (ODE)

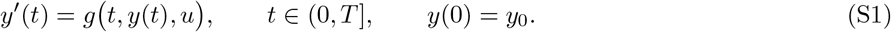

In all three benchmarks, the control enters one component of *g* through a regularized bolus input *I*(*t, u*). For administration times

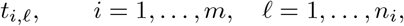

and a short infusion window *ε >* 0, the implemented input is the piecewise-constant regularization

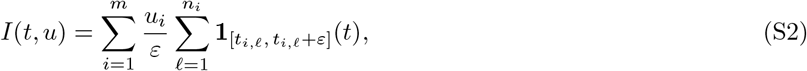

or, in some cases, a scalar dose *u* ∈ ℝ applied at *M* fixed times,

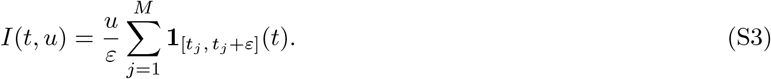

The indicator function **1**_*A*_ equals 1 on the set *A* and 0 otherwise.

##### Cost functional and reduced problem

Each test model specifies an observable quantity

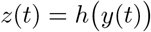

(e.g. a biomarker or tumor volume) and a reference trajectory *z*_ref_(*t*). The cost functional is of the form

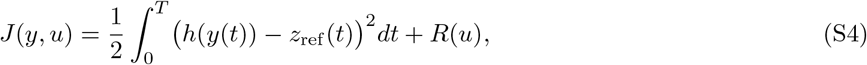

where *R* is the direct dose penalty. A quadratic penalty takes the form 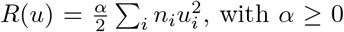, with *α* ≥ 0 and *n*_*i*_ administrations sharing. The IDR and BiTE benchmarks use *R* = 0; TGI uses the linear dose penalty stated below.

The *reduced* cost is defined by substituting the solution of Eq. (S1) into the cost functional,

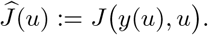

The dose optimization problem solved by OptiDose is

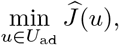

where *U*_ad_ ⊂ ℝ^*m*^ encodes box constraints on the doses. In the present derivations we ignore these constraints and focus on first-order optimality.

##### Adjoint and dose gradient

For a running loss *ℓ* and no terminal loss, define the adjoint ***λ***(*t*) by

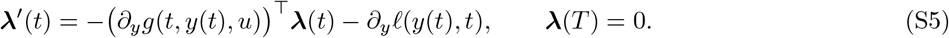

If the dosing input enters state component *j*_0_, the reduced gradient is

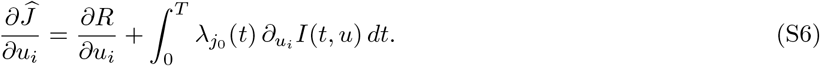

The direct penalty derivative ∂*R/*∂*u*_*i*_ is zero for IDR and BiTE and *α* = 10^−7^ for TGI. For the width-*ε* regularized bolus in Eq. (S2), this becomes

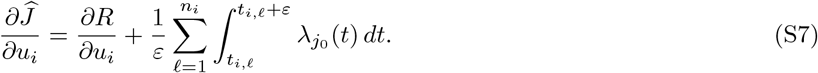

The same adjoint definition is used in Supplementary Note 5.

##### Indirect response (IDR) biomarker model

###### Model equations

The IDR test model describes a PK compartment *C*(*t*) and a biomarker *B*(*t*):

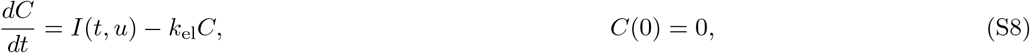

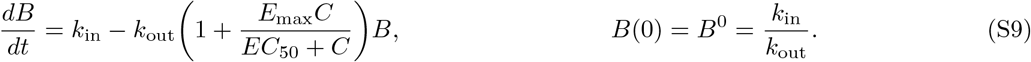

The state vector is *y* = (*C, B*)^⊤^ ∈ ℝ^2^, so *n* = 2. The control *u* contains *m* dose amplitudes; in the OptiDose benchmark *m* = 6 weekly cycles with *n*_*i*_ = 7 daily applications per week.

###### Reference trajectory and loss

The observable is the biomarker,

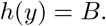

The reference trajectory *B*_ref_ is specified piecewise as a quadratic decay from the baseline *B*^0^ to a target value *B*_tar_ over 7*m*_1_ days (with *m*_1_ = 2 in the benchmark) and then held constant:

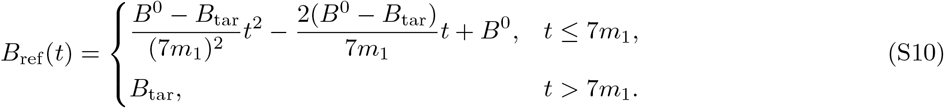

The cost functional (S4) becomes

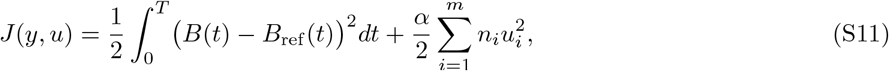

with *T* = 42 days and, in the benchmark, *α* = 0.

###### Jacobian of the right-hand side

Writing Eqs. (S8)–(S9) as *y*^′^ = *g*(*t, y, u*), the Jacobian ∂_*y*_*g* ∈ ℝ^2×2^ is

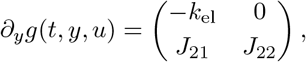

where

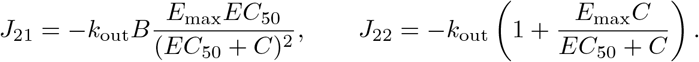

###### Adjoint equation

The running loss is

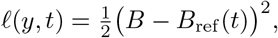

so

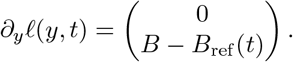

Let ***λ*** = (*λ*_*C*_, *λ*_*B*_)^⊤^. Inserting the Jacobian and ∂_*y*_*ℓ* into Eq. (S5) yields

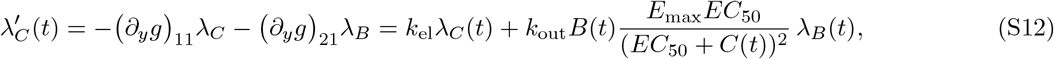

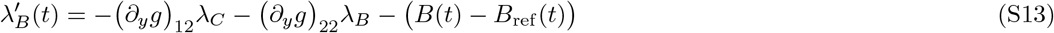

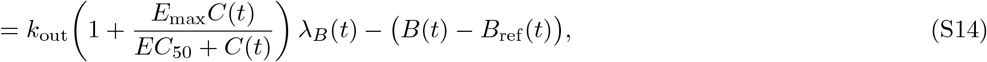

with terminal condition *λ*_*C*_(*T*) = *λ*_*B*_(*T*) = 0.

###### Dose gradient

In the IDR model, the input *I*(*t, u*) acts directly on *C*, so *j*_0_ is the first state component and 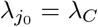. Combining Eqs. (S2) and (S6) gives

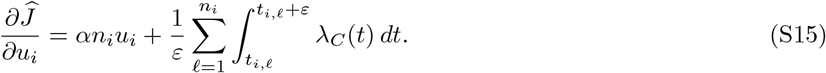

This is the analytic gradient formula implemented and compared against automatic differentiation a nd finite differences in the JAX code.

##### Tumor growth inhibition (TGI) model

###### Model equations

The TGI test model couples an oral PK model with a nonlinear tumor growth and three damage compartments. The state vector is

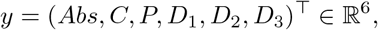

where *Abs* is the absorption compartment, *C* the central concentration, *P* the proliferating tumor volume and *D*_1_, *D*_2_, *D*_3_ are damage compartments. Let

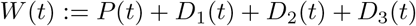

denote the total tumor burden. The PKPD model is

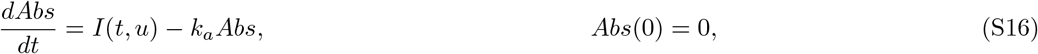

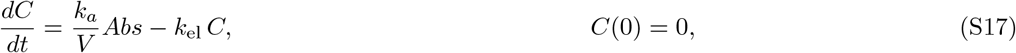

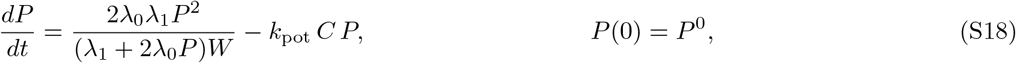

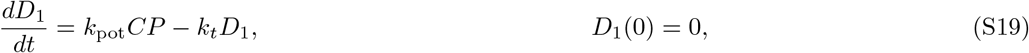

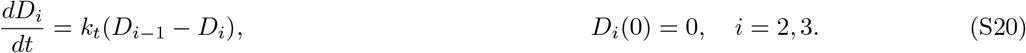

The parameter vector is

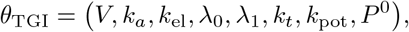

whose numerical values were retained unchanged from the published OptiDose benchmark.^1^

###### Reference trajectory and loss

The observable is the total tumor burden,

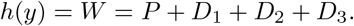

The reference trajectory used in the tracking objective is

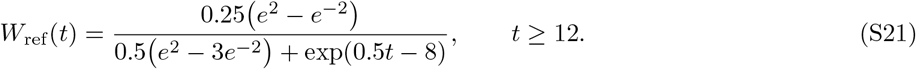

Only times from day 12 onward contribute to the tracking term, so no extension of *W*_ref_ to [0, 12) is required. The cost functional is

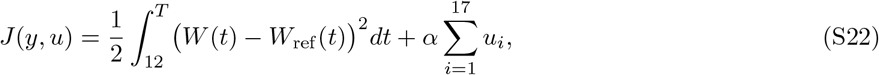

with *T* = 30 days and *α* = 10^−7^ in the benchmark setup. The 17 controls are independent oral-dose amplitudes *u*_*i*_ administered once daily at *t*_*i*_ ∈ {12, 13, …, 28} days.

###### Jacobian

In the state order (*Abs, C, P, D*_1_, *D*_2_, *D*_3_), rows 1–2 describe PK, row 3 describes tumor growth, and rows 4–6 describe the damage chain. The analytic Jacobian is

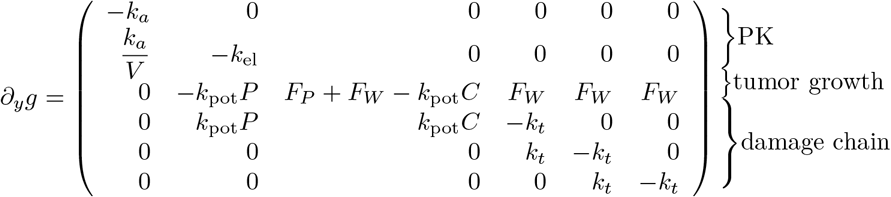

Here

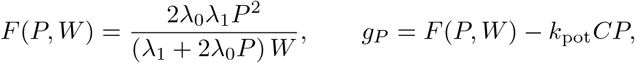

and, with the other argument held fixed, define

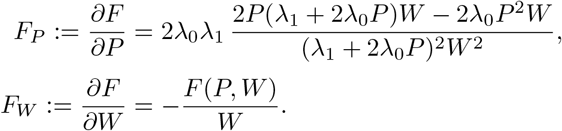

As *W* = *P* + *D*_1_ + *D*_2_ + *D*_3_, the *P*-row contains *F*_*P*_ + *F*_*W*_ in its *P* column and *F*_*W*_ in each damage-state column.

###### Adjoint equation

The running loss is

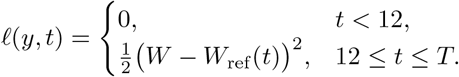

Its state gradient is zero for *t <* 12. For 12 ≤ *t* ≤ *T*, since *W* depends only on *P, D*_1_, *D*_2_, *D*_3_, we have

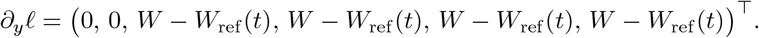

For 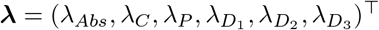, the adjoint is given by Eq. (S5) with terminal condition ***λ***(*T*) = 0.

###### Dose gradient

In the TGI model, the dosing input acts on the absorption compartment, so *j*_0_ corresponds to *Abs* and 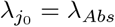. Combining the linear dose penalty with the regularized bolus in Eq. (S2) gives

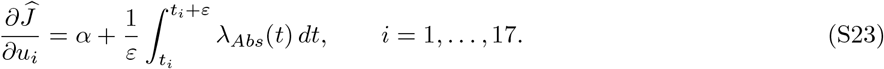

##### Bispecific T cell engager (BiTE) model

###### Model equations

The BiTE benchmark represents a bispecific antibody with target-mediated drug-disposition dynamics. The labels *A* and *B* distinguish the two binding partners: *R*_*A*_ and *R*_*B*_ denote their free receptors, *RC*_*A*_ and *RC*_*B*_ the corresponding binary complexes, and *RC*_*AB*_ the ternary complex. The remaining states are central concentration *C*, peripheral amount *AP*, and absorption-compartment amount *Abs*. The state vector is

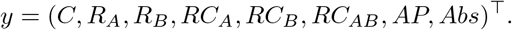

The ODE system is

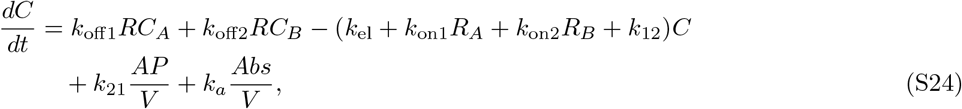

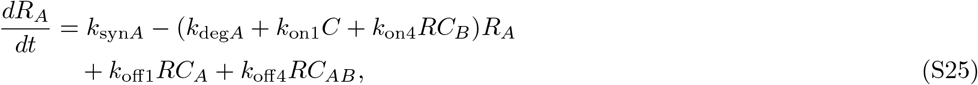

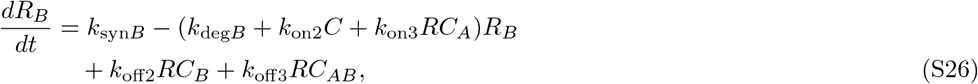

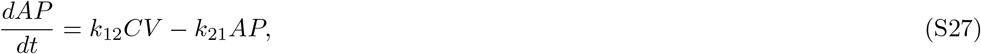

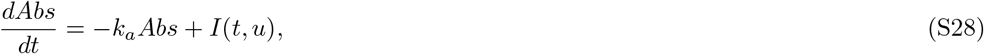

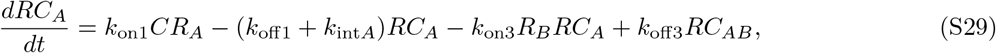

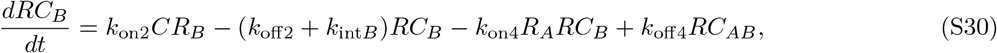

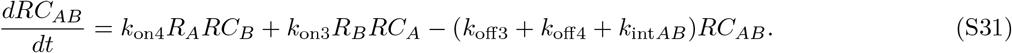

Initial conditions are

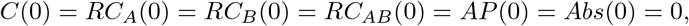

while the free receptors start at their steady–state values,

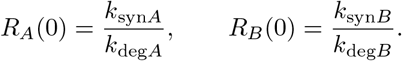

The parameter values below are those used in the OptiDose benchmark and retained unchanged in the replication:

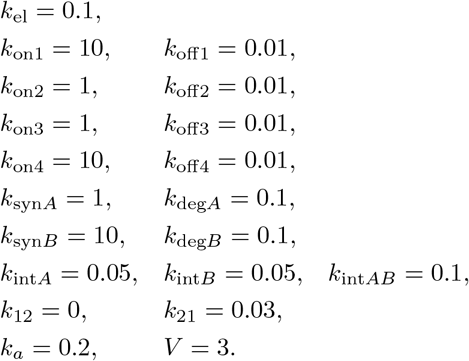

###### Reference value and loss

The observable quantity in this benchmark is the ternary complex

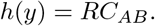

The reference value is the minimum of the two receptor steady states,

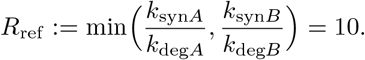

The cost functional is

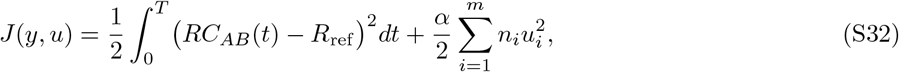

with *T* = 140 days and *α* = 0 in the benchmark.

###### Jacobian

Let *y* = (*y*_1_, …, *y*_8_)^⊤^ denote the state vector in the order

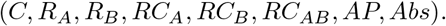

The Jacobian in this state order is

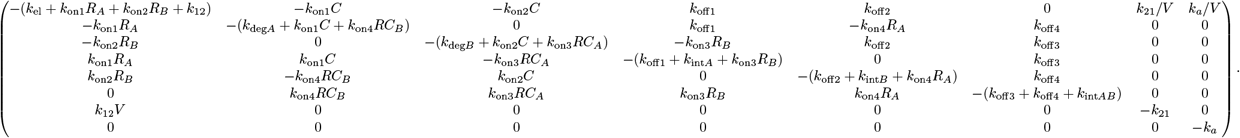

###### Adjoint equation

The running loss is

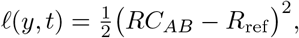

so

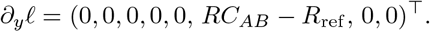

For ***λ*** = (*λ*_*C*_, *λ*_*R*_, *λ*_*R*_, *λ*_*RC*_, *λ*_*RC*_, *λ*_*RC*_, *λ*_*AP*_, *λ*_*Abs*_)^⊤^, the adjoint is given by Eq. (S5) with terminal condition ***λ***(*T*) = 0.

###### Dose gradient

This benchmark assigns one shared scalar amplitude *u* to all three administrations rather than optimizing three independent dose amplitudes. Because *I*(*t, u*) enters only the equation for the absorption compartment, the corresponding adjoint component is *λ*_*Abs*_, associated with the eighth state (*j*_0_ = 8). For regularized boluses as in Eq. (S3) at *t*_*j*_ ∈ {0, 48, 96} with window length *ε*, we have

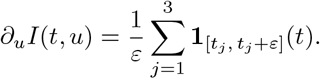

The reduced cost 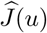 therefore satisfies

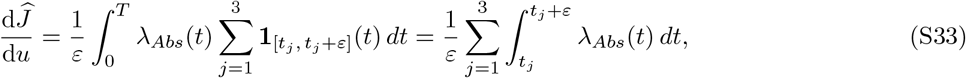

with *α* = 0 in this benchmark. If one instead treats the three doses as independent controls *u*_1_, *u*_2_, *u*_3_, the formula generalises to

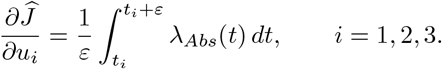

### Supplementary Note 2: Differentiable relaxation of quantized dose controls

#### Gumbel–Softmax relaxation

##### Quantized dose formulation

For *m* fixed administration times, let **u** = (*u*_1_, …, *u*_*m*_) denote the dose-amplitude vector and let ℒ (**u**) be the reduced objective obtained by simulating the mechanistic model. If each dose must be an integer multiple of a fixed increment *a >* 0, the admissible levels for event *j* are

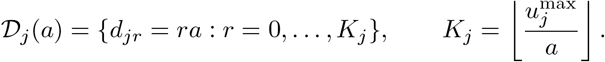

The exact quantized problem is

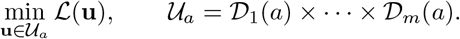

##### Differentiable categorical relaxation

The optimization variables are per-event logits 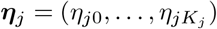, rather than the dose amplitudes themselves. For independent *g*_*jr*_ ~ Gumbel(0, 1), the relaxed categorical probabilities are

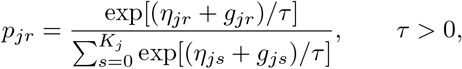

and the probability-weighted dose supplied to the simulator is

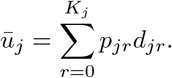

Because 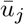 is a convex combination of admissible grid points, it satisfies the event-wise box bounds by construction. If 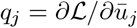, then, conditional on the sampled Gumbel variables,

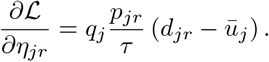

A feasible quantized regimen is recovered by maximum-probability projection,

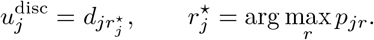

This construction permits gradient-based optimization of categorical probabilities while enforcing the clinical dose grid after projection.^2,3^ The same formulation can quantize administration times by replacing the dose grid *D*_*j*_(*a*) with a finite set of clinically admissible administration times. Projection then returns a feasible discrete time while the relaxed control remains differentiable during optimization.

### Supplementary Note 3: Neutropenia DDE validation and implementation

#### Timing-gradient validation

The simulated repeated-daily comparator administered filgrastim on days 1–13 after each chemotherapy dose. Clinical guidelines and CHOP-14 studies describe repeated daily administration of short-acting filgrastim^4,5^. Prior analyses of this model showed that the timing of granulocyte colony-stimulating factor (G-CSF) administration can strongly and non-intuitively affect the depth and duration of neutropenia^6^. DiffDose therefore parameterized each cycle directly by one filgrastim administration time. The smoothed dosing map described below places these seven timing controls in the continuous DDE right-hand side, allowing forward-mode AD to differentiate the implemented objective with respect to administration time.

Implementing the Craig *et al*. model in a differentiable program required validating both the modified simulator and the gradients used for optimization. The smoothed dosing map and AD-compatible delay handling make dose time differentiable but change the numerical representation of the original impulsive treatment inputs. We therefore confirmed the expected absolute-neutrophil count (ANC) dynamics and compared the seven-component AD gradient with central finite differences of the complete objective evaluation.

ForwardDiff gradients agreed with central finite differences using *h* = 10^−4^ day. The relative *ℓ*_2_ error between the seven-component gradient vectors was 2.9 × 10^−5^, the maximum component-wise absolute error was 3.6 × 10^−6^, and the directional derivatives agreed to a relative error of 1.4 × 10^−4^. These checks support the numerical reliability of the AD gradients for the smoothed, discretized DDE objective.

#### Model definition

The timing case study used the 12-state granulopoiesis and chemotherapy-induced neutropenia model, with corresponding code translated from MATLAB^6^. The model couples hematopoietic stem-cell and neutrophil maturation dynamics to two-compartment G-CSF pharmacokinetics and four-compartment chemotherapy pharmacokinetics. The equations below state the mathematical model first, followed by the treatment inputs and numerical implementation. The published biological structure and source parameterization were retained; our study-specific changes were the Julia translation, differentiable treatment inputs used to optimize administration times, and the numerical and optimization workflow described below.

##### States and delays

**Supplementary Table 4.** States in the implemented neutropenia DDE.

| State | Interpretation |
| --- | --- |
| $Q$ | Hematopoietic stem-cell pool |
| $N$ | Circulating neutrophils; $ANC = 8.19N$ |
| $G_1$ | Free plasma G-CSF |
| $\tau_{NM}$ | State-dependent post-mitotic maturation time |
| $C_1$ | Central chemotherapy compartment |
| $A_Q$ | Stem-cell amplification factor |
| $A_N$ | Neutrophil-lineage amplification factor |
| $C_2$ | First peripheral chemotherapy compartment |
| $C_3$ | Second peripheral chemotherapy compartment |
| $N_R$ | Marrow neutrophil reservoir |
| $C_4$ | Third peripheral chemotherapy compartment |
| $G_2$ | Bound or tissue-associated G-CSF |

For any state *X*, define

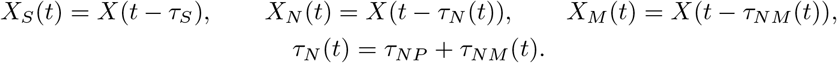

Here, *τ*_*S*_ is the fixed stem-cell cycling delay, *τ*_*NP*_ is the fixed precursor proliferation time, and *τ*_*NM*_ (*t*) is the state-dependent maturation time. Thus *t* − *τ*_*N*_ (*t*) + *τ*_*NP*_ = *t* − *τ*_*NM*_ (*t*), whereas *X*_*N*_ and *X*_*M*_ generally refer to different delayed times.

##### Complete 12-state system

Let *R*_*C*_(*t*) denote chemotherapy input and *I*_*G*_(*t*) exogenous G-CSF input. States and inputs without explicit time arguments are evaluated at *t*. The system is

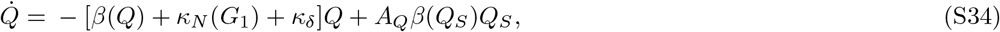

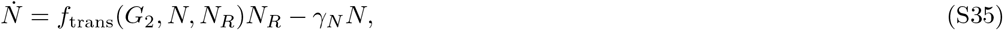

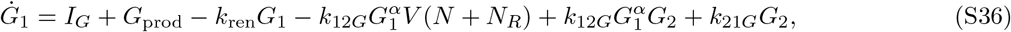

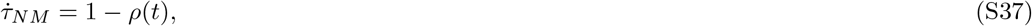

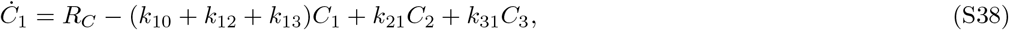

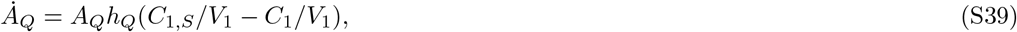

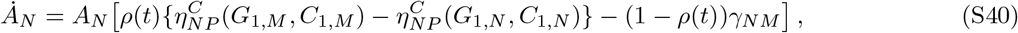

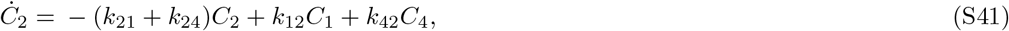

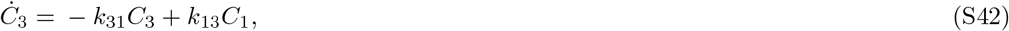

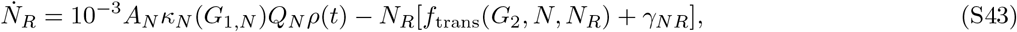

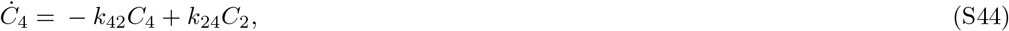

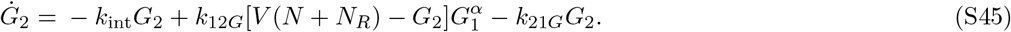

The dimensionless maturation-velocity ratio is

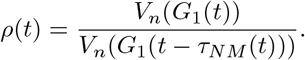

It compares the current maturation velocity with its value at the delayed time and enters both the maturation-delay equation and the delayed cell flux. The auxiliary functions describe stem-cell self-renewal *β*, G-CSF-regulated differentiation *κ*_*N*_, precursor proliferation 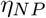, chemotherapy-suppressed proliferation 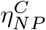, maturation velocity *V*_*n*_, and reservoir release *f*_trans_:

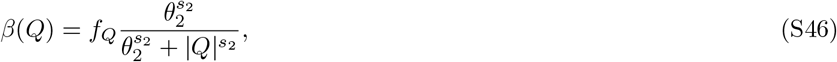

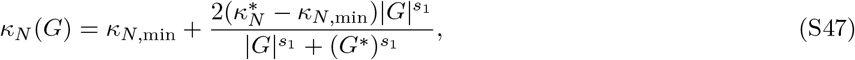

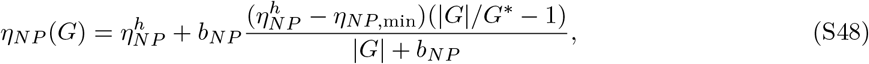

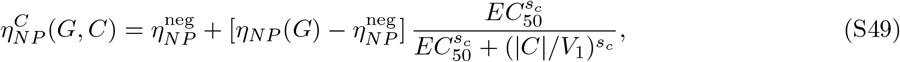

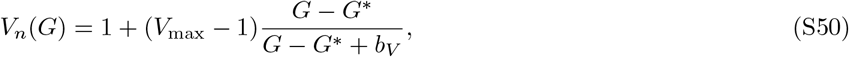

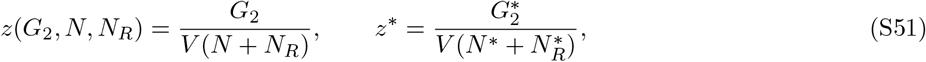

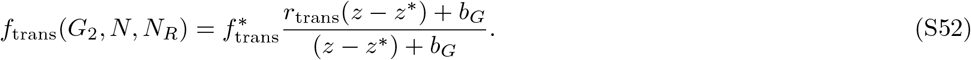

The factor 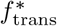 preserves the homeostatic reservoir-release rate. The factor 10^−3^ in Eq. (S43) is the unit-scaling coefficient retained from the published implementation.^6^

##### Homeostatic history and initial conditions

The history is constant for *t* ≤ 0 and equal to the homeostatic initial vector in Table 5. Derived entries are recomputed from the base parameters whenever a scenario is constructed.

**Supplementary Table 5.**
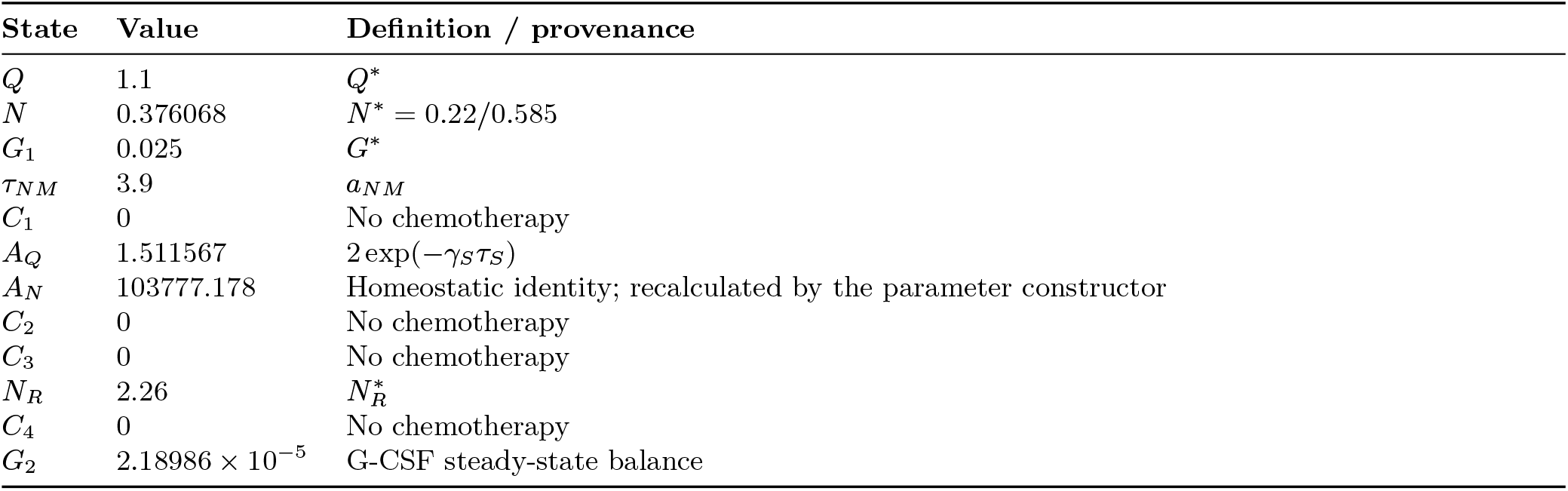
Initial history for the 12-state DDE.

| State | Value | Definition / provenance |
| --- | --- | --- |
| $Q$ | 1.1 | $Q^*$ |
| $N$ | 0.376068 | $N^* = 0.22/0.585$ |
| $G_1$ | 0.025 | $G^*$ |
| $\tau_{NM}$ | 3.9 | $a_{NM}$ |
| $C_1$ | 0 | No chemotherapy |
| $A_Q$ | 1.511567 | $2 \exp(-\gamma_S \tau_S)$ |
| $A_N$ | 103777.178 | Homeostatic identity; recalculated by the parameter constructor |
| $C_2$ | 0 | No chemotherapy |
| $C_3$ | 0 | No chemotherapy |
| $N_R$ | 2.26 | $N_R^*$ |
| $C_4$ | 0 | No chemotherapy |
| $G_2$ | $2.18986 \times 10^{-5}$ | G-CSF steady-state balance |

##### Fixed, estimated, and calculated parameters

The implementation distinguishes three parameter roles. *Fixed* values are constants carried from the published model or its MATLAB implementation; *estimated* values are parameters fitted in the source-model calibration; and *homeostatically calculated* values are determined from steady-state identities and recalculated by the Julia constructor whenever a scenario is created^6^. Exact values, formulas, code-level names, equations, dosing logic, benchmark configuration, parity tests, and provenance are available in the DiffDose repository. Units follow the source model.

##### Differentiable inputs, objective, and numerical solution

Seven chemotherapy administrations occurred at 14-day intervals over *T* = 98 days. Each chemotherapy amount was 4, 000 mg m^−2^ × 1.723 = 6, 892 in the central-compartment input and was delivered over 1*/*24 day. One 300 *µ*g filgrastim administration time was optimized independently within each cycle. Relative times were constrained to [0.5, 13.0]^7^ days and initialized at day 4.

For chemotherapy starts *c*_*j*_ = 14(*j* − 1) days, *j* = 1, …, 7, duration Δ_*C*_, and total central-compartment amount *D*_*C*_, the benchmark used

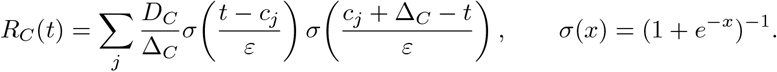

The product of the two sigmoid factors approximates a rectangular chemotherapy infusion while providing differentiable onset and offset times. For a filgrastim injection of amount *D*_*G*_ at *g*_*j*_ = *c*_*j*_ + *w*_*j*_, where *w*_*j*_ is the optimized cycle-relative time in days, the input was

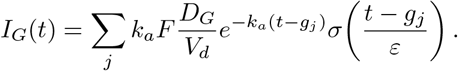

Here the sigmoid gates the start of the usual first-order subcutaneous absorption profile. The smooth width was *ε* = 5 × 10^−4^ days. This representation places the administration times inside the continuous right-hand side so that their dual-number partials propagate through the numerical solve.

The equations above state the published model separately from a numerical safeguard introduced in our Julia translation. In code, the absolute numerical floor *ε*_AD_ = 10^−30^ is added before fractional powers, (|*x*| + *ε*_AD_)^*ν*^, where *ν* denotes the applicable exponent, including the G-CSF binding exponent *α* and chemotherapy Hill exponent *s*_*c*_. This floor is not a solver tolerance. It avoids undefined dual-number partials at zero while remaining negligible at physiologically relevant concentrations; it is not part of the source mathematical model.

The objective was the cumulative ANC deficit

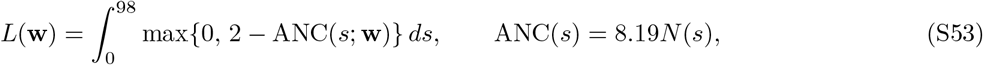

This is the main-text ANC-deficit objective evaluated with ANC expressed numerically in 10^9^ cells/L; the physical threshold of 2 × 10^9^ cells/L is therefore 2, and losses are reported in 10^9^ cells L^−1^ day. A custom composite trapezoidal rule evaluated the integral on states saved every 0.05 day.

State-dependent delays were evaluated inline and were not registered through `dependent_lags`. The model was integrated with `MethodOfSteps(RK4())`, reltol = abstol = 10^−8^, and *dt*_max_ = 0.01 day.^7^ ForwardDiff propagated seven-component dual numbers through this solver and the trapezoidal objective:^8^

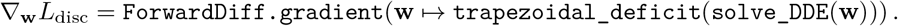

Accordingly, *L*_disc_ denotes the *smoothed, discretized objective*; these are its AD gradients, not analytic sensitivities of the original state-dependent DDE with impulsive treatment inputs. Central differences with step 10^−4^ day were used as an independent numerical check. Continuous forward sensitivities and continuous adjoints were not used because the available workflow did not provide the required state-dependent DDE sensitivity equations and delay/event terms. The tested reverse-mode solver configurations were also impractical for the long, tightly stepped trajectory because their tape, trajectory, or recomputation requirements exhausted available memory.

The AD gradients were supplied either to adaptive moment estimation (Adam)^9^ (learning rate = 0.05, *β*_1_ = 0.9, *β*_2_ = 0.999) or to bounded L-BFGS through `Fminbox(LBFGS())`.^10,11^ This optimizer comparison is downstream of the differentiation route: both methods consumed gradients of the same smoothed discretized objective.

**Supplementary Table 6.**
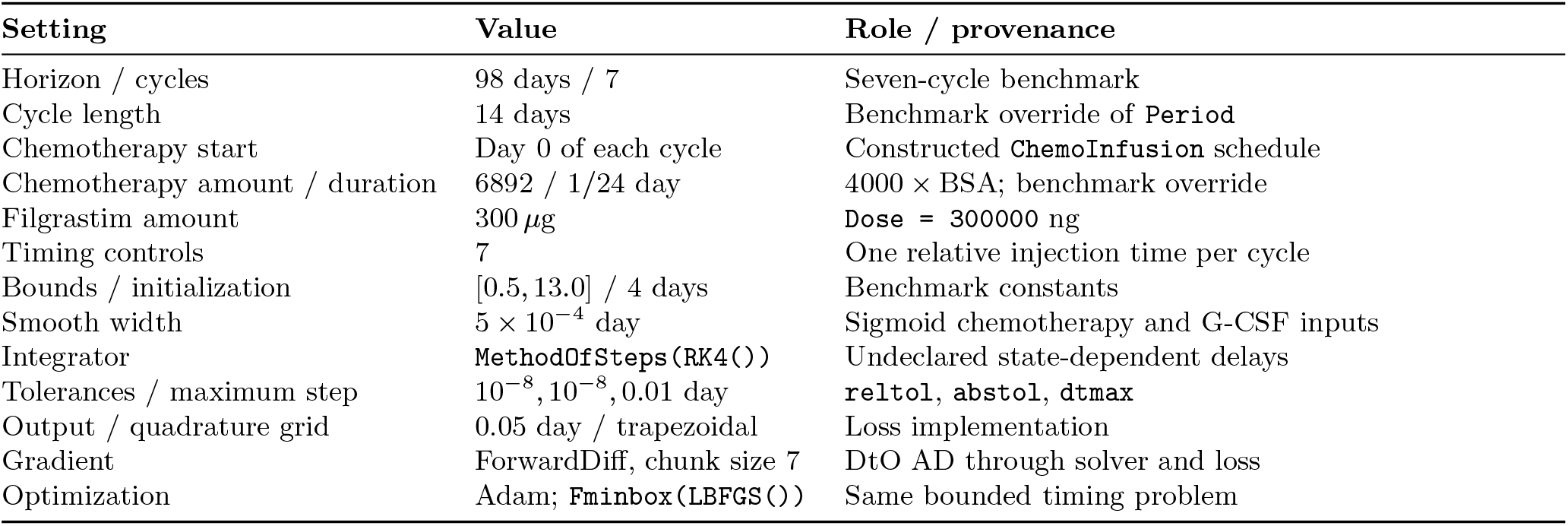
Treatment and numerical settings for the seven-cycle timing benchmark.

##### Declared-delay compatibility patch

During development, we also tested forward-mode AD with the state-dependent lags registered through `dependent_lags`. In the archived `DelayDiffEq.jl` 5.61.0 environment, the discontinuity tracker passed the output of `discontinuity_function` to a scalar bracketing solver and subsequently stored the located discontinuity time in a Float64 queue. When a timing control was seeded by `ForwardDiff`, this function could instead return a `ForwardDiff.Dual`, producing a type error inside `discontinuity_time` before the gradient calculation completed.

For this compatibility test, we evaluated the following transformation of the root-function value:

~~~
val = discontinuity_function(integrator, lag, T, t + theta * dt)
val isa ForwardDiff.Dual ? ForwardDiff.value(val) : val
~~~

The transformation supplied the bracketing solver with the primal root value while leaving dual-number propagation active in the DDE right-hand side and state integration. Because it removes derivatives of the internally located discontinuity time itself, we treat this as an implementation compatibility experiment rather than a general sensitivity rule for state-dependent DDEs. The seven-cycle results reported here were generated with the unpatched, undeclared-lag route described above. The focused compatibility example and analysis implementation are available in the DiffDose repository; the example documents why declared-lag tracking failed in the original AD workflow and the workaround that was evaluated.

### Supplementary Note 4: Mosunetuzumab virtual population construction and optimization

#### Virtual population validation

The independently constructed 250-member virtual population (VPop) reproduced the qualitative behavior reported by Hosseini *et al*. across four published step-up regimens, including IL-6, activated T cell, tumor-response, and responder-fraction summaries (Supplementary Figure 7). Supplementary Figures 8 and 9 then compare fixed and individualized regimens and show the convergence of each patient’s optimization.

**Supplementary Figure 7.**
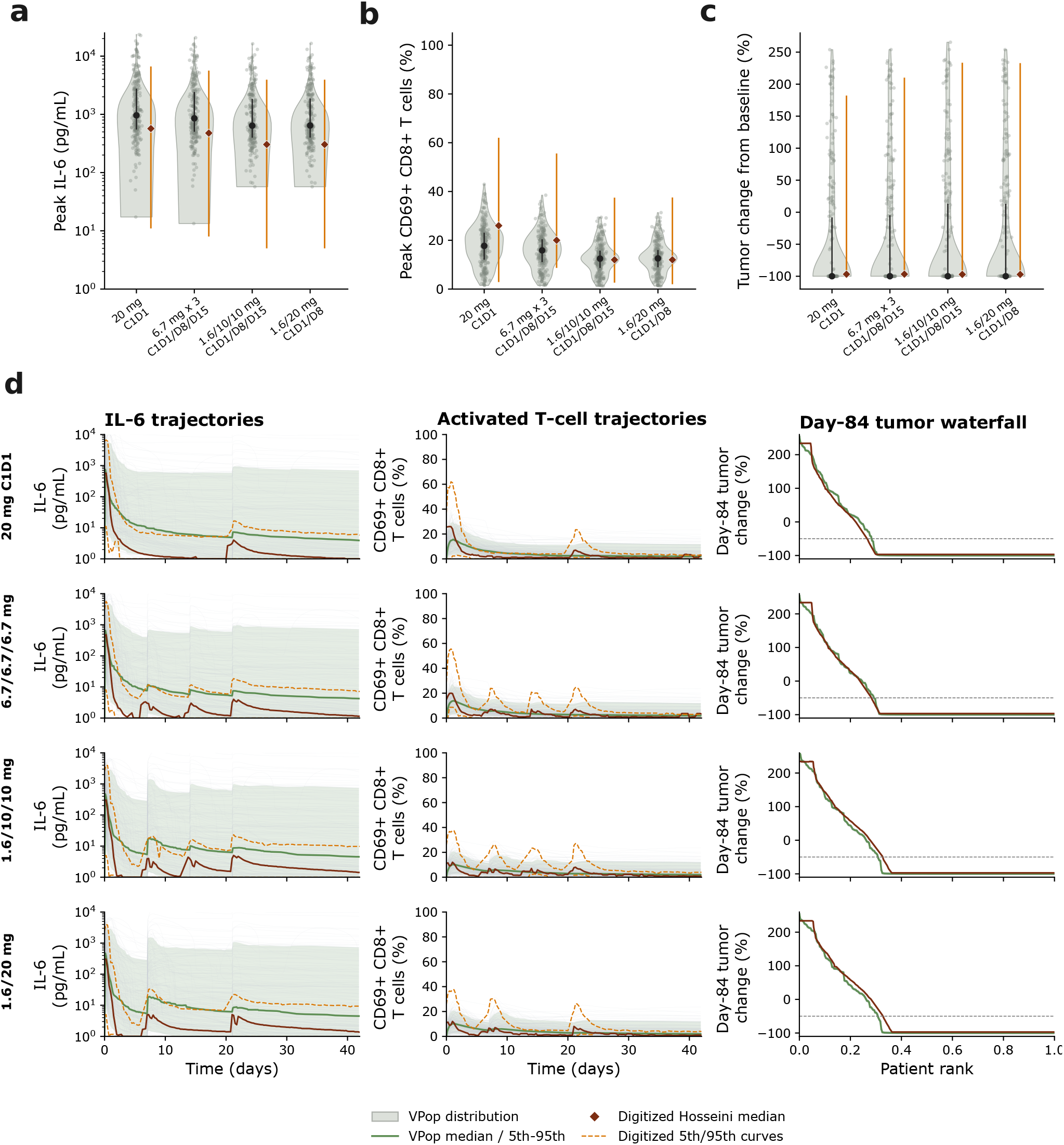
The independently constructed virtual population reproduces the published mosunetuzumab response summaries. (a–c) Distributions of peak IL-6, peak peripheral CD69^+^CD8^+^ T cells, and day-84 tumor change across four step-up regimens. Gray-green violins show the 250-member VPop; black markers and bars show its median and interquartile range; orange markers and intervals show values digitized from Hosseini et al. (d) Corresponding trajectory and tumor-waterfall comparisons. Thin gray curves show virtual patients, green curves and bands show the VPop median and 5th–95th percentiles, and orange curves show digitized published summaries. The dashed horizontal line marks 50% tumor reduction.

**Supplementary Figure 8.**
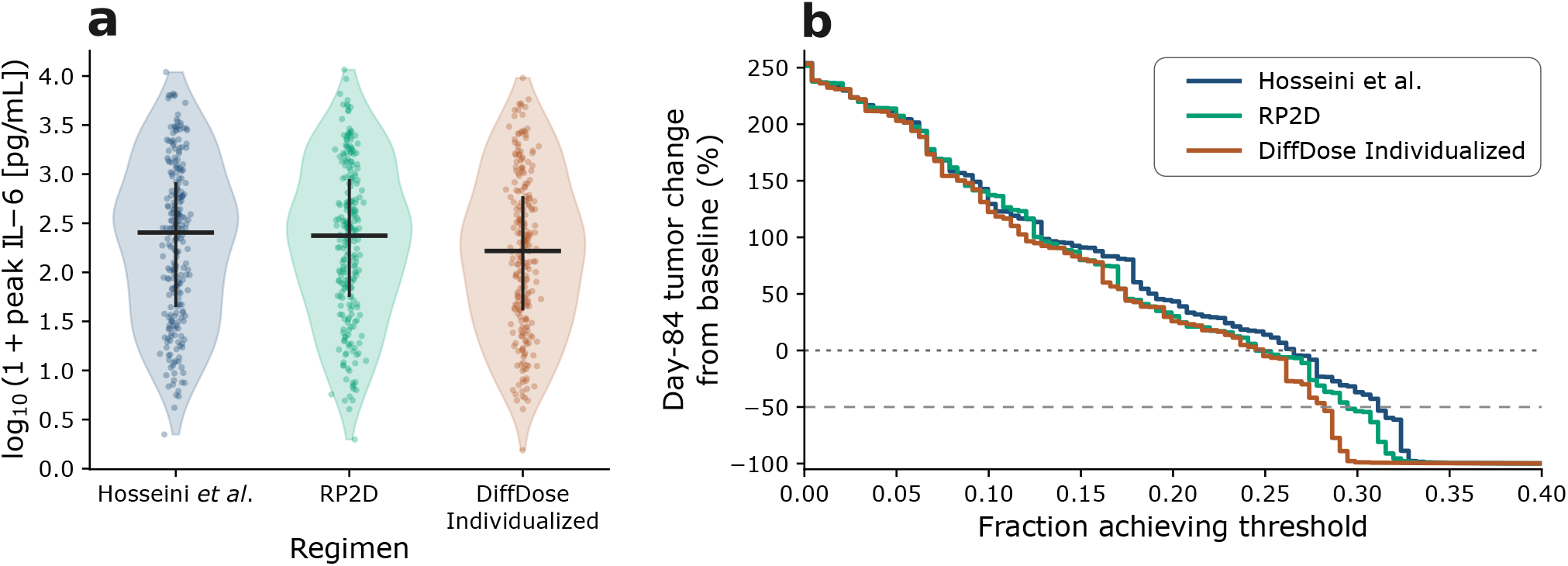
Endpoint distributions under fixed and individualized mosunetuzumab regimens. (a) Global IL-6 peak over 84 days. Violin plots show the population distributions, points show individual virtual patients, and black markers and bars show the median and interquartile range. (b) Reverse empirical cumulative distributions of day-84 tumor change. The horizontal reference lines mark no change from baseline and 50% tumor reduction. Curves compare the Hosseini et al. schedule, the recommended phase 2 dose (RP2D), and individualized DiffDose regimens.

**Supplementary Figure 9.**
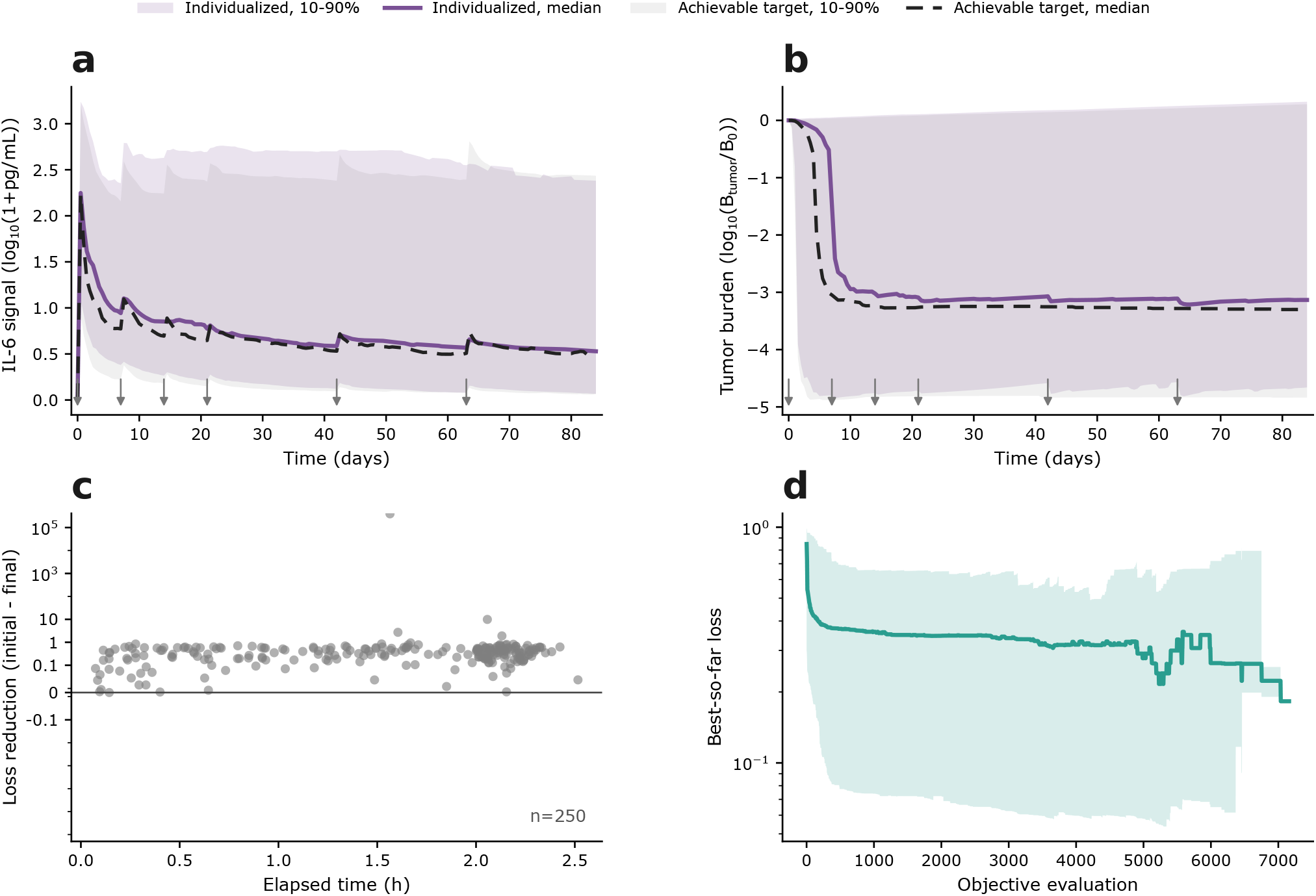
Individualized objective construction and optimization histories. (a,b) Median optimized IL-6 and tumor trajectories with 10th–90th percentile population bands, overlaid on the patient-specific achievable targets. The tumor target came from each patient’s fixed 60-mg trajectory, and the IL-6 target came from that patient’s fixed 1-mg trajectory. Arrows mark administration days. (c) Per-patient loss reduction versus elapsed optimization time; positive values indicate improvement from the initialized dose vector. (d) Median best-so-far loss with a 10th–90th percentile band across objective evaluations.

#### Mechanistic model and fixed simulation protocol

The mosunetuzumab quantitative systems pharmacology model was translated from the SimBiology models reported by Hosseini *et al*. and Susilo *et al*. into the typed Julia implementation MosunModelCore.^12,13^ The production model contains 36 dynamic states and preserves the published model structure and nominal parameterization, including mosunetuzumab disposition and target binding, T cell activation and trafficking, B cell killing and tumor dynamics, and IL-6 production and disposition. Our additions were the typed Julia handling of repeated assignments and bolus events, construction of an independent virtual population, and the patient-specific objective and optimization workflow. Repeated-assignment quantities required for tumor, T cell, B cell, drug, and IL-6 endpoints were reconstructed after each state evaluation. Mosunetuzumab dose events target the central amount state and are converted from milligrams to *µ*g kg^−1^ using a fixed 70-kg body weight.

Each simulation covered 84 days. The stiff ODE was solved with Rodas4P, abstol = 10^−8^, reltol = 10^−5^, and output every 0.5 day. Fixed dose times were included as solver stops, and the proposed post-event time step was 0.01 day. The six fixed administration days represented three doses in cycle 1 (days 0, 7, and 14) followed by one dose in each of cycles 2–4 (days 21, 42, and 63):

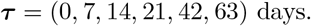

Tumor burden, the global IL-6 peak, tumor and IL-6 areas under the curve, total dose, and central drug exposure were computed from the saved trajectory. The objective used for optimization included only day-84 tumor burden, global IL-6 peak, and the 20-mg cycle-dose target; the remaining endpoints were post hoc comparisons.

##### Virtual population construction and released parameter files

The candidate population was generated using the extended Fourier amplitude sensitivity test (eFAST) over 62 inputs spanning diffuse large B-cell lymphoma (DLBCL)-range model parameters, curated cytokine-response variables, latent tumor-burden variables, and three inert dummy-null variables. Positive parameters designated for log-scale sampling were sampled in log_10_ space. Parameters with stable total-effect signals above the dummy-null thresholds, together with curated tumor- and IL-6-response variables, defined 33 high-importance parameters.^14^ Candidates were screened against digitized Hosseini *et al*. IL-6, activated T cell, day-84 tumor-response, and responder-fraction summaries across the four fixed regimens. We then resimulated 20,000 prefiltered candidates and selected a 250-member subset by stochastic pruning that balanced agreement with these four outcome summaries while penalizing collapse of high-importance parameter variability. The selection search used seed 20260505.

**Supplementary Table 7.** Category-level summary of the 62-parameter VPop sampling universe.

| Parameter family | Count | Applied | Sampling role |
| --- | --- | --- | --- |
| Other mechanistic model parameters | 28 | 28 | DLBCL-range model variability |
| T cell activation and infiltration | 14 | 14 | Immune activation and trafficking |
| PK and exposure | 5 | 5 | Mosunetuzumab disposition |
| Tumor burden, growth, and infiltration | 5 | 3 | Baseline and tumor dynamics; two latent selection variables |
| B cell killing | 4 | 4 | Drug-mediated tumor-cell killing |
| IL-6 | 3 | 3 | Cytokine production and disposition |
| Dummy-null controls | 3 | 0 | Empirical sensitivity thresholds |
| Total | 62 | 57 | 38 log-scale and 24 linear sampling specifications |

Supplementary Data (`Supplementary_Data.csv`) gives the exact 250 selected parameter vectors together with their candidate and eFAST provenance identifiers. The complete 62-parameter sampling specification, including lower and upper bounds, base values, sampling scales, source classes, parameter families, and implementation notes, is versioned in the public DiffDose repository.

##### Independent six-dose objective for each virtual patient

For virtual patient *i*, the six independent controls were

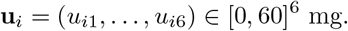

Two fixed simulations provided achievable, patient-specific references. The schedule **r**_max_ = (60, 60, 60, 60, 60, 60) mg provided the day-84 tumor target, whereas **r**_min_ = (1, 1, 1, 1, 1, 1) mg provided the global IL-6-peak target. Define the smooth approximation to the positive-part function as

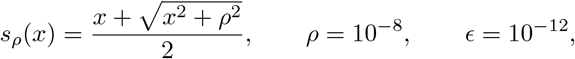

where *ρ* controls the smoothing scale and *ϵ* prevents division by zero. Define the normalized residual-tumor measure *R*_*i*_ and global IL-6-peak measure *P*_*i*_ as

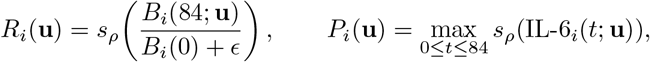

Here, *B*_*i*_ is the tumor B-cell state, and IL-6_*i*_ combines plasma IL-6 with the model’s volume-weighted tissue and tumor contributions. The peak is evaluated over the saved trajectory. The targets are

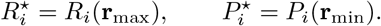

The one-sided, normalized tumor term was

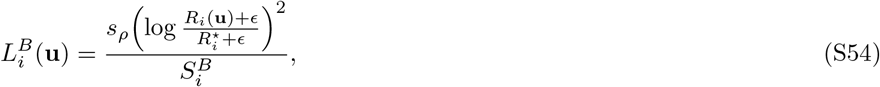

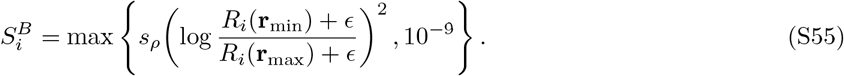

The IL-6 term, in the same orientation, was

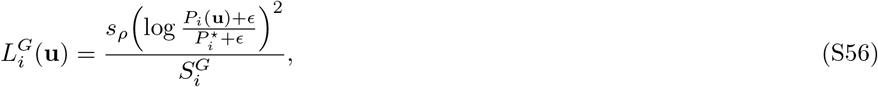

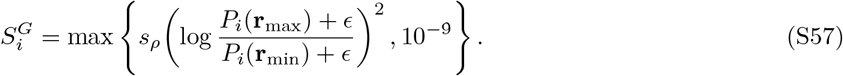

Schedules better than the corresponding target therefore received no material additional reward. Cycle totals were

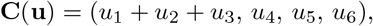

and the dose-target term was

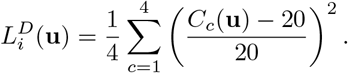

The implemented per-patient problem was

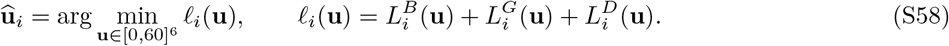

with all three weights set to one. The reported individualized analysis solved Eq. (S58) separately for each of the 250 virtual patients; it did not minimize an expectation over the cohort.

For comparison, a cohort-shared extension would optimize one regimen for all virtual patients,

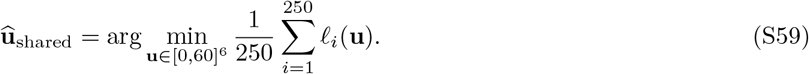

This mean-loss problem was considered as an exploratory objective family, but it was not used for the individualized results reported in the main manuscript.

##### Differentiation and bounded optimization

The dose amplitudes were propagated through fixed-time dose events and the `Rodas4P` solve with `SensitivityADPassThrough()`. `ForwardDiff.gradient!` was used to differentiate the discretized simulation and endpoint calculation directly. The resulting six-component gradient was used as input to `Fminbox(BFGS())`.^10,11^ Each virtual patient was initialized independently from Uniform(0, 60)^6^ with seed 20260519. Limits were 50 BFGS iterations, 1000 objective calls, and 7200 seconds per patient. From this, we retained the dose vector with the lowest logged finite objective value for each patient.

**Supplementary Table 8.** Code-aligned settings for individualized virtual population optimization.

| Component | Setting | Implementation |
| --- | --- | --- |
| Cohort | $n = 250$ | Supplementary Data |
| Controls | Six doses in $[0, 60]$ mg | Days 0, 7, 14, 21, 42, 63 |
| Tumor target | Six 60-mg doses | Patient-specific day-84 ratio |
| IL-6 target | Six 1-mg doses | Patient-specific global peak |
| Dose target | 20 mg per cycle | Cycle 1 sum; cycles 2–4 single doses |
| Integrator | `Rodas4P` | 84 days; 0.5-day saved output |
| Tolerances | $10^{-8}$ absolute; $10^{-5}$ relative | Fixed across patients |
| Gradient | `ForwardDiff.gradient!` | DtO through solver and endpoints |
| Optimizer | `Fminbox(BFGS())` | Box-constrained quasi-Newton update |
| Initialization | `Uniform [0, 60]`<sup>6</sup>, seed 20260519 | Independent for each patient |

##### Exploratory objective families

Before fixing the endpoint-constrained objective in Eq. (S58), we evaluated three exploratory families. Reference-based step-up objectives compared four grouped controls with either the Hosseini double-step regimen or the recommended phase 2 dose schedule. Safety-constrained objectives penalized increases in IL-6 peak and IL-6 AUC while allowing tumor benefit only in the safety-preserving region. Trajectory objectives compared the full tumor and IL-6 time courses with patient-specific high-dose and low-dose targets. These variants were used to select an interpretable primary formulation; they were not pooled into an expected cohort loss and were not used for the objective reported in the main individualized analysis. The cohort-shared objective in Eq. (S59) and neural-network policy objectives were not used to generate any reported result; they are retained to define extensions discussed in the main Discussion.

### Supplementary Note 5: Differentiation routes and downstream optimization

#### First-order derivatives for finite-dimensional dose control

The calculations in this study apply established differentiation tools to dose-regimen optimization; they do not introduce a new automatic-differentiation algorithm. Consider a fixed schedule 0 *t*_1_ *<* ≤ · · ·*< t*_*m*_ *< T*, one dose control per event, **u** = (*u*_1_, …, *u*_*m*_)^⊤^, and a state **x**(*t*) satisfying

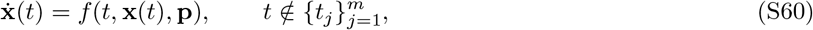

with fixed-time reset maps

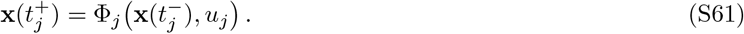

For the scalar-dose map, *D*_*uj*_ Φ_*j*_ is an *n*-vector. When the map is written as a function of the full control vector **u**, its control Jacobian is *n* × *m* and acts on *δ***u**; the scalar-dose derivative acts on *δu*_*j*_. The pre-dose initial state **x**(0^−^) = **x**_0_ is fixed; a dose at time zero is included through the reset map. The reduced objective is

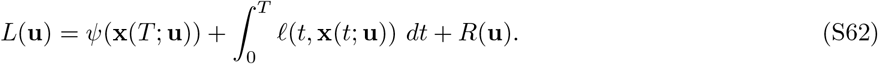

Here, *ψ* is the terminal loss, *ℓ* the running loss, and *R* the direct dose penalty. The trajectory functional ℱ in the main text comprises the terminal and integrated running-loss terms. Gradients are column vectors. The central numerical task is to evaluate ∇_**u**_*L* and supply it to a finite-dimensional optimizer.

#### Equivalent variational, Lagrange-multiplier, and PMP equations

For a perturbation *δ***u**, the state variation satisfies the tangent equation

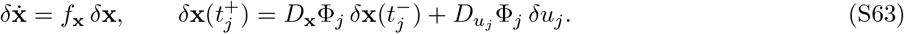

With *δ***x**(0^−^) = 0, these variations give the trajectory contribution to the dose gradient; the direct contribution is ∇_**u**_*R*. Alternatively, introduce a time-dependent Lagrange multiplier ***λ***(*t*), integrate the constrained first variation by parts, and choose ***λ*** to eliminate the unknown state variation. This gives

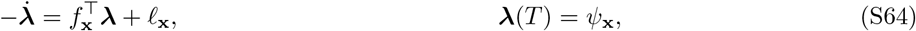

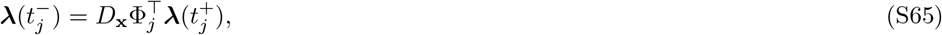

and the componentwise gradient

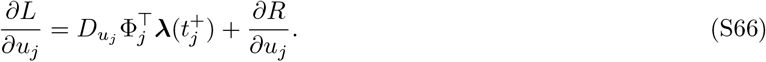

For an additive bolus into compartment 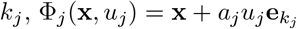, where *a*_*j*_ is a fixed dose-to-state conversion coefficient. The adjoint is continuous across the event and

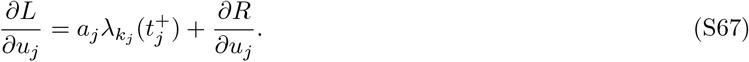

Variational calculus, the Lagrange-multiplier construction, and Pontryagin’s maximum principle (PMP) therefore produce the same first-order tangent, costate, and stationarity equations, up to sign and Hamiltonian conventions. They are alternative derivations of the same reduced gradient, not different gradient algorithms. Equations (S63) and (S64) describe the continuous derivative problem; automatic differentiation can instead differentiate the numerical program that approximates Eqs. (S60)–(S62). Because the event times above are fixed, no event-time term is required. State-, parameter-, or control-dependent event times require additional hybrid sensitivity or saltation terms.

#### Differentiation routes

The forward/reverse distinction specifies how derivative information is accumulated, whereas the optimize-then-discretize/discretize-then-optimize distinction specifies whether the continuous equations or the implemented solver is differentiated. These choices are independent of the optimizer that later consumes the gradient.^15–17^ Continuous forward sensitivities propagate tangent states, whereas continuous adjoints solve a costate backward after the state trajectory. Discrete forward AD propagates Jacobian–vector products through the executed solver operations, and discrete reverse AD propagates vector–Jacobian products with stored states, checkpointing, or recomputation. We evaluated all four routes in the ODE benchmarks. In the DiffDose applications described in the main text, we used direct forward AD in all three case studies and direct reverse AD as an additional ODE benchmark variant.

Finite differences perturb the controls and repeatedly evaluate the discretized objective. They provide a useful independent gradient check but are neither a forward nor reverse AD route, and their cost and accuracy depend on the differencing step and solver tolerances. Where just-in-time compilation was used, benchmark timings were measured after a warm-up call so that one-time compilation did not dominate route comparisons.

#### Downstream optimizers and software

Automatic differentiation, continuous sensitivities, adjoints, and finite differences determine the gradient supplied to an optimizer; they do not determine the dose update. Adam is a first-order adaptive method,^9^ whereas BFGS and L-BFGS use successive gradients to build curvature approximations; L-BFGS-B or a bounded wrapper enforces box constraints.^10^ Nelder–Mead and simulated annealing were used as derivative-free comparisons.^18,19^ Initialization, scaling, bounds, and stopping rules remain consequential, and none of these local methods certifies a global optimum for a nonconvex mechanistic objective. Comparable MATLAB interfaces include `fminunc, fmincon`, and `lsqnonlin`.

The ODE benchmarks used JAX/Diffrax^20^ with SciPy L-BFGS-B. The Julia case studies used the SciML ecosystem: the neutropenia model used `DelayDiffEq.jl`^7^ and `ForwardDiff.jl`^8^ with Adam or bounded L-BFGS, whereas the mosunetuzumab VPop used `ForwardDiff.jl` with bounded BFGS in `Optim.jl`.^11^ Within each route comparison, the forward model, objective, bounds, initialization, and downstream optimizer were held fixed. Case-specific solver versions, tolerances, event representations, and optimizer settings are given in Supplementary Notes 1, 3, and 4.

